# Phospholipid Scramblases TMEM16F and Xkr8 mediate distinct features of Phosphatidylserine (PS) externalization and immune suppression to promote tumor growth

**DOI:** 10.1101/2025.04.17.649445

**Authors:** Varsha Gadiyar, Rachael Pulica, Ahmed Aquib, James A. Tranos, Christopher Varsanyi, Trevor Frederick, Ziren Wang, Luis Fernandez Almansa, Lawrence Gaspers, Mariana S De Lorenzo, Sergei V Kotenko, Sushil Tripathi, Roger W. Howell, Alok Choudhary, David C. Calianese, Raymond B. Birge

## Abstract

The phospholipid scramblases Xkr8 and TMEM16F externalize phosphatidylserine (PS) by distinct mechanisms. Xkr8, is activated by caspase-mediated proteolytic cleavage, and in synergy with inactivation of P4-ATPase flippases, results in the irreversible externalization of PS on apoptotic cells and an “eat-me” signal for efferocytosis. In contrast, TMEM16F is a calcium activated scramblase that reversibly externalizes PS on viable cells via the transient increase in intracellular calcium in live cells. The tumor microenvironment (TME) is abundant with exposed PS, resulting from prolonged oncogenic and metabolic stresses and high apoptotic indexes of tumors. Such chronic PS externalization in the TME has been linked to host immune evasion from interactions of PS with inhibitory PS receptors such as TAM and TIM receptors. Here, in an effort to better understand the contributions of apoptotic vs live cell PS-externalization to tumorigenesis and immune evasion, we employed an E0771 orthotopic breast cancer model and genetically ablated Xkr8 and TMEM16F using CRISPR/Cas9. While neither the knockout of Xkr8 nor TMEM16F showed defects in cell intrinsic properties related to proliferation, tumor-sphere formation, and growth factor signaling, both knockouts suppressed tumorigenicity in immune-competent mice, but not in NOD/SCID or RAG-KO immune-deficient strains. Mechanistically, Xkr8-KO tumors suppressed macrophage-mediated efferocytosis, and TMEM16F-KO suppressed ER stress/calcium-induced PS externalization. Our data support an emerging idea in immune-oncology that constitutive PS externalization, mediated by scramblase dysregulation on tumor cells, supports immune evasion in the tumor microenvironment. This links apoptosis/efferocytosis and oncogenic stress involving calcium dysregulation, contributing to PS-mediated immune escape and cancer progression.

## Introduction

The composition of lipid modalities and their distribution in the plasma membrane lipid bilayer is critical for maintaining the biochemical and biophysical properties of the membrane ^1, 2^. In eukaryotic cells, plasma membrane phospholipids are asymmetrically distributed across the membrane whereby phosphatidylcholine (PC) and sphingomyelin (SM) are mostly comprised to the outer leaflet while phosphatidylethanolamine (PE), phosphatidylinositol (PI) and phosphatidylserine (PS) are mostly restricted to the inner leaflet ^3^. In resting cells at homeostasis, the negative anionic charge of PS is also important for the recruitment of cytosolic signaling and anchoring proteins with polybasic regions as well as regulating phospholipid exchange reactions between the plasma membrane and the endoplasmic reticulum via lipid transfer proteins ^4–8^. The maintenance of PS asymmetry in healthy cells is primarily regulated by P4-ATPase and related flippases that actively facilitate the vectorial transfer of PS from the outside to the inner membrane such that, under homeostasis, the PS is almost completely restricted to the intracellular leaflet ^9–11^.

However, cells can rapidly disrupt PS and phospholipid asymmetry under a variety of physiological conditions and expose PS and PE to the outer cell surface by lipid scramblases. Emblematically, during apoptosis, caspase mediated activation of the lipid scramblase Xkr8 (and concomitant inactivation of P4 ATPase flippases) results in irreversible PS externalization in the dying cells and signal for the apoptotic cells to the internalized and degraded by efferocytosis ^12–14^. By contrast, during cell activation or the transient rise in intracellular calcium, cells can activate the calcium-dependent TMEM16 family of lipid scramblases ^15, 16^, to transiently and reversibly externalizing PS. In both cases, by redirecting PS from the inside of the membrane to the outer membrane, these lipid scramblases effectively switch PS from having an intracellular itinerary of PS-binding proteins (i.e the intracellular PS proteome) to having an extracellular itinerary of PS-binding proteins (i.e the extracellular PS proteome). On PS-positive apoptotic cells, externalized PS interacts with a host of inhibitory PS receptors, including TIMs, TAMs, CD36, and BAI1 that promote efferocytosis ^17^ ^18–20^ and tolerance ^21–23^ ^24, 25^.On activated cells, such as T cells, NK cells, dendritic cells (DCs), and macrophages, externalized PS can interact with immune resolving factors that dampen inflammation to prevent collateral tissue damage ^26–28^. Moreover, on platelets and endothelial cells, externalized PS can recruit coagulation factors that facilitate fibrin formation and wound repair ^29^. Such PS-interacting proteins convey many physiological and homeostatic regulatory functions to maintain tissue function.

In contrast to the physiological and homeostatic events associated with PS externalization under = conditions described above, PS is constitutively externalized in the tumor microenvironment by mechanisms that are still not completely understood ^18, 19^. Many highly proliferative tumors have high apoptotic indexes that necessitate excessive efferocytosis that have been correlated as cold tumors with poor survival prognosis and outcomes ^30, 31^. In addition, most solid tumors are associated with metabolically stressed and hypoxic vasculature, which externalize PS, preempting the development of PS-targeting monoclonal antibodies (mAbs) such as Bavituximab ^32–36^. PS externalization in the tumor microenvironment (TME) has also been associated with exhausted T cells, Tregs, and myeloid suppressor cells ^34, 35, 37^, as well as the polarization of macrophages towards wound healing M2 phenotypes as well as maintaining immature DCs ^38^. Although many solid tumors display constitutive PS externalization that can be assessed by honing of PS-targeting antibodies ^39–41^ and Annexin V^42^ ^43^, the mechanisms of PS externalization in the tumor microenvironment are surprisingly not well understood, nor are the composition of PS-positive cells in the TME well articulated.

Here, we investigated the contributions of two phospholipid scramblases, Xkr8 and TMEM16F, that are implicated in PS externalization in the context of immune regulation. We hypothesize that tumor cells are a major source for externalized PS in the TME. Using an E0771 luminal B orthotopic breast cancer model, we show that E0771 cells preferentially express both Xkr8 and TMEM16F over other isoforms, and that E0771 tumor-bearing transplanted mice externalized PS in the tumor microenvironments as evident by the homing of PS-targeting mAbs. Subsequently, using CRISPR/Cas9 gene editing to individually knockout Xkr8 or TMEM16F and, we show that both PS scramblases directly contribute to immune regulation and tumor growth in immune-competent mice but not NOD/SCID or RAG immune-deficient mice. Our data support a complex and duality mechanism for PS externalization in the tumor microenvironment whereby apoptosis/efferocytosis of growing tumors and calcium-stressed PS-out viable tumor cells both likely contribute to PS-mediated immune escape and tumor progression.

## Methods

### Cell culture

All cells except Expi293 were cultured in a humidified incubator at 37° C and 5% CO_2_. EO771 and HEK293 cells were cultured in DMEM ([+] 4.5 g/L glucose, L-glutamine [-] sodium pyruvate) supplemented with 10% FBS and 1% Penicillin/Streptomycin in 100 mm cell culture treated dishes. MC38 and B16 F-10 cells were cultured in DMEM ([+] 4.5 g/L glucose, L-glutamine [-] sodium pyruvate) supplemented with 10% FBS, 1% Penicillin/Streptomycin and additional 5% L-Glutamine Streptomycin in 100 mm cell culture treated dishes. Jurkat cells, W3 - I1dm and W3 - CDC50A^ED29^ cells (suspension cells) were cultured in RPMI supplemented with 10% FBS and 1% Penicillin/Streptomycin in T75 culture flasks. TAM reporter CHO (Mertk-γ-R1 and EGFR/TAM) cell lines were cultured in HAMS F-12 media supplemented with 10% FBS, 1% Penicillin/Streptomycin and additional 5% L-Glutamine Streptomycin in 100 mm cell culture treated dishes. BMDMs (Bone marrow derived macrophages) were cultured in IMDM supplemented with 10% Heat Inactivated FBS, 1% Penicillin/Streptomycin and 20 ng/mL of recombinant mouse M-CSF. Expi293 cells were cultured in Expi293 expression media in Erlenmeyer flasks in a humidified incubator at 37° C and 8% CO_2_ on a shaker at 110 rpm. Cell treatments were done for 24 hours with 10 μM MG132, or glucose free DMEM for glucose deprivation.

### BMDM isolation

Bone marrow derived macrophages were isolated from the tibia and femurs of 8-12 weeks old C57BL/6 mice. The tibia and femurs were isolated and washed for 15 seconds each, once in 70% ethanol and then in PBS, following which their ends were cut to reveal the bone marrow. The bones were placed cut end facing down in a 0.5 mL Eppendorf tube with a hole pierced with an 18 G needle, which was placed in a 1.5 mL Eppendorf tube containing 80 μL of BMDM media. The tubes were then centrifuged at 15,000 g to allow the bone marrow to precipitate into the 1.5 mL Eppendorf tube. The cell pellet was treated with 5 mL of RBC lysis buffer for 5 minutes at room temperature, later quenching with BMDM media. The cells were collected by centrifuging at 300 g for 5 minutes and filtered using a 70-micron cell strainer. The cells were counted using trypan blue exclusion using a hemocytometer and were plated in 150 mm cell culture treated plates at 10^7^ viable cells/mL, in a total volume of 30 mL. Recombinant murine MCSF (Biolegend) was added at a final concentration of 20 ng/mL. On day 3 of differentiation, the 15 mL of the cell culture medium was removed and replaced with 15 mL of fresh MCSF containing medium. BMDMs were fully differentiated on day 7. Differentiated BMDMs were detached gently using a scraper and re-plated into 6 well, 12 well or 24 well plates based on experimental needs.

### Expression Constructs

The protein sequences for Bavituximab (US8956616B2), 1N11 (US20180289771A1), and 11.31 (US20110318360A1) were retrieved from their respective patents, and the corresponding DNA sequences are obtained through reverse translation using the Expasy Translate tool. Gene fragments for the VH (variable heavy) and VL (variable light) chains of these antibodies are designed with overlapping regions of approximately 18 bp to ensure compatibility with the respective backbone vectors, and are synthesized by Genewiz (Azenta Life Sciences). The pVITRO1-M80-F2-human IgG1/κ and pVITRO1-dV-humanIgG1/λ plasmids are isolated from 5 mL bacterial cultures using the Qiagen Miniprep kit, with the backbone regions amplified by PCR using Q5 high-fidelity polymerase. The PCR products are analyzed by agarose gel electrophoresis and purified with a PCR purification kit (Qiagen). The NEB Builder HiFi assembly cloning kit is used to ligate the four DNA fragments (backbone regions and variable chains), and the resulting products are transformed into E. coli (DH5-alpha), with colonies selected on LB agar plates containing Hygromycin. At least 16 colonies are selected, and colony PCR is performed to identify positive hits, followed by plasmid purification and Sanger sequencing to confirm the correct clone. The pCP-CMV-GCaMP6f plasmid was purchased from Addgene (#40755). The plasmid was transfected into WT and TMEM16F KO cells using a Lipofectamine LTX kit. G418 was used to select for successfully transfected clones (600 μg/ml).

### Protein expression in Expi293 system

Expi293 cells were cultured in suspension at 37°C with 8% CO2 on an orbital shaker, maintained at a density of 1-2 × 10LJ cells/mL. Plasmids for the antibody constructs were purified using the Qiagen Endtotoxin free Maxiprep kit and assessed for quality and quantity by Nanodrop. On the day of transfection, the cells were counted by trypan blue exclusion and diluted to a density of ∼1.5 × 10LJ cells/mL with >95% viability using Expi293 expression media. 100 μg of the plasmid DNA and 300 μL of the Expifectamine reagent were diluted in 5 mL of Opti-MEM media in separate tubes. The dilutions were incubated for 5 minutes at room temperature. The transfection complex was formed by mixing plasmid DNA with Expifectamine reagent in Opti-MEM media and incubated for 15 minutes at room temperature. The transfection complex was added to a 100 mL culture of cells by pipetting slowly and swirling the flask. The cells were incubated for 18 hours, after which transfection enhancers were added. For Gla containing proteins (ProS, Gas6, Gas6-IFN fusion proteins), Vitamin K was added at a final concentration of 2μg/mL or Warfarin was added at a final concentration of 2mM to produce γ-Carboxylated (Active) or non-γ-Carboxylated (Inactive) versions of the proteins respectively. The cell culture was incubated for an additional 72 hours to allow antibody secretion into the supernatant, which was then collected. The cells were collected by centrifugation at 2000 rpm for 10 minutes and the supernatant was passed through a 0.2 micron filter to remove debris. The protein containing supernatant was then incubated with 1 mL of TALON beads (for His tagged proteins) or 1 mL of Protein-A agarose beads (for antibodies) respectively. The supernatant-affinity bead mixture was incubated for 48 hours at 4°C.

### Protein purification

After incubating the protein supernatant with beads for 48 hours, the mixture was centrifuged to collect the beads. The supernatant was then passed through a gravity-flow protein purification column containing the beads, with the flow-through collected. For antibody purification, the column was washed with PBS, and elution buffer (Glycine pH 2.5) was added to elute the antibody in 500 µL fractions, which were neutralized with Tris-HCl buffer and kept on ice to prevent precipitation. The fractions were pooled and concentrated using centrifugal concentrators, then dialyzed overnight in PBS. For His tagged proteins, the column was washed with 250 mM Phosphate buffer pH 8, 1.5 M NaCl pH8, and the protein was eluted with 200 mM Imidazole diluted in the phosphate buffer. The protein concentration was determined by a Bradford assay, with a yield of 1-2 mg of antibody per 100 mL culture. The purified proteins were aliquoted and kept at −80°C for long term storage.

### Antibody and protein labelling

Antibodies were fluorescently labelled using the Dylight 680 or the Dylight 800 NHS ester labelling kit. The Dylight 680 Maleimide reactive kit was used to label any protein containing the Gla domain. To prepare the protein for labeling, a 0.05M sodium borate buffer at pH 8.5 was used. The protein was dissolved in phosphate-buffered saline (PBS) to avoid precipitation, as buffers containing primary amines like Tris or glycine could interfere with the labeling reaction. For each labeling reaction, 100-500µL of purified protein at 1-2.5mg/mL was used, and if necessary, a buffer exchange into the labeling buffer was performed via centrifugal concentrators. For the labeling reaction, the protein solution was transferred to the DyLight 680 NHS-Ester dye, mixed well, and incubated at room temperature for 1 hour. After incubation, excess dye was removed using Thermo Scientific Dye Removal Columns. It was ensured that unreacted dye was completely removed to achieve optimal results. The labeled protein was stored at 4°C, protected from light, for up to one month, or in single-use aliquots at −20°C for long-term storage. To calculate degree of labelling, A280 and the absorbance at A680 were measured using Nanodrop.

### Zr-89 labelling and PET/CT imaging

Conjugation of 11.31 and Isotype antibodies with ^89^Zr was done in accordance with Vosjan and colleagues ^44^. Briefly, p-isothiocyanatobenzyl-desferrioxamine (Df–Bz–NCS) was conjugated to lysine groups of the mAbs at pH 9, followed by purification using PD-10 columns ^44^. [^89^Zr]-oxalic acid solution was mixed with the Df-Bz-NCS-mAb and incubated at room temperature for 1 hour. Unlabeled ^89^Zr was removed by purifying the conjugated antibody using a PD-10 column. The labelling efficiency of the reaction was determined by ITLC. To conduct PET/CT imaging, approximately 200 μg of [^89^Zr]-Df-Bz-NCS-mAb in 200 μL solution was injected intra-peritoneally into each mouse before PET/CT imaging with an MILabs VECTor^6^ (Utrecht, Netherlands).

### Scramblase Knockouts using CRISPR/Cas9 gene editing

EO771 TMEM16F and Xkr8 KO cells were developed by Synthego using the RNP method. Clones were made by sorting single cells into a 96 well plate using FACS Aria II. MC38 TMEM16F KO cells were created by designing sgRNAs using Genscript. sgRNAs and Cas9-EGFP were co-transfected into MC38 cells using the CRISPR MAX transfection kit. 48 hours after transfection, GFP was seen in transfected cells and were sorted into single cell clones by FACS Aria II. Clones were screened using Western blot, surveyor assay and q-RT-PCR to validate the knock-out.

Ano6-2-TGGAAATAAGAGTGGACGCG

Ano6-3-TGACACTCTCGTTCACCCGG

Ano-5-CCTTCTCGTAGATCGTGTTG

Xkr8- 1- CGTTGAACCAGGAGAAATAG

Xkr8- 3- CGTACTGGACAACGGCCCAC

Xkr8- 7- CGACCACGTCTAAGGCCACA

### Murine Cancer models

8-12 weeks old female C57BL/6 mice (000664) or NSG mice (NOD.Cg-*Prkdc^scid^ Il2rg^tm1Wjl^*/SzJ, 005557) were purchased from Jackson Laboratories. The Rag1 KO mice (C.129S7(B6)-*Rag1^tm1Mom^*/J, 003145) were a gift from Bergsbaken Lab (CII, Rutgers University). Homozygous pairs were mated to produce mice for the experiments. Mouse experiments were performed in accordance with the guidelines and under the approval from Institutional Animal Care and Use Committee at the Rutgers University, Rutgers New Jersey Medical School (Newark, NJ). The mice were housed in pathogen-free facility, maintained under a strict 12-hour light cycle, and given regular chow diet.

EO771, or MC38 cells were counted and resuspended in DMEM serum free media at 10^6^ cells/ 100 μL. The mice were anesthetized using ketamine/xylazine and the left inguinal mammary gland or the flank were depilated using Nair. 50 μL of the cell suspension was mixed with 50 μL of Matrigel and 100 μL of the cell-Matrigel suspension was injected into the mammary gland fat pad or the flank. The tumor volume was measured every 3 days using an electronic caliper and the tumor volumes were calculated using the formula: Length of the tumor x (Width of the tumor)^2^ x 0.5263. The mice were sacrificed when the tumor reached 2000 mm^3^, after which the tumors were dissected, and the tumor weights and spleen weights were measured. IVIS imaging was conducted using the IVIS Spectrum. Mice bearing EO771 tumors of ∼200-500 mm^3^ were injected intraperitoneally with NIR dye labelled PS targeting antibodies and fluorescent imaging was done after 24, and 48 hours after injection. In vivo calcium imaging was done by injecting EO771 cells expressing the GCAMP6F plasmid and waiting for the tumors to reach ∼200-500 mm^3^. Mice were fed Alfalfa free chow (LabDiet 5V75) to minimize background GFP signal in the body.

### Flow cytometry

Cells were collected by trypsinization for adherent cells and by centrifugation for suspension cells. Cells were resuspended and counted by trypan blue exclusion. 1 × 10^6^ cells were collected in a V bottom 96 well plate or a 1.5 mL Eppendorf tube and washed with cell staining buffer (PBS + 1% BSA). Fc block was diluted to 1:50 dilution and 50 μL added to cells and mix by pipetting. Cells were incubated at 4°C for 20 minutes. Primary antibody cocktail was prepared and 50 μL was added to each sample. Cells were incubated 4°C for 30 minutes protected from light. After primary antibody staining, cells were washed with 100 μL cell staining buffer, and then 100 μL of diluted secondary antibody added and pipetted to mix. Cells were incubated at 4°C for 30 minutes protected from light, after which the cells were washed with 100 μL cell staining buffer. For fixation, cells were incubated with 4% paraformaldehyde diluted in cell staining buffer for 15 minutes at room temperature protected from light. Wherever applicable, cells were permeated using BD Fix perm buffer and stained using active caspase 3 antibody for 30 minutes. Finally, cells were washed with cell staining buffer and resuspend in 300 μL of cell staining buffer and transfer to BD flow tubes. Samples were acquired using BD Fortessa or BD LSR II. Data analysis was done with Flow Jo software. For annexin V staining, the BD Apoptosis detection assay kit was used and the given protocol was followed. Cytosolic calcium staining was performed in cell culture by treating with 3 μM Fluo-4-AM for 30 minutes at 37 C in cell culture and cells were collected and processed for Annexin V staining as mentioned.

### Western blotting

Standard SDS-PAGE and wet transfer methods were used for Western blot. Briefly, protein containing supernatants or cell lysates were diluted in 6X sample loading buffer and subjected to SDS PAGE (either 8, 10 or 12 % resolving gel and a 4 % stacking gel) electrophoresis to separate the proteins, and further transferred onto a PVDF membrane. The blots were incubated in blocking buffer 5% non-fat dry milk for 1 hour at room temperature, after which they were incubated with the primary antibody β-Actin, p-Akt (CST), TMEM16F(Courtesy Lily Jan, UCSF), Xkr8 (Abclonal) at 1:1000 dilution at 4°C overnight. The blots were washed thrice with TBST for 15 minutes each, after which they were incubated with an appropriate secondary-HRP antibody (1:4000 or 1:10,000). The blots were washed thrice in TBST for 15 minutes each, after which they were developed using ECL substrate and imaged using Biorad GelDoc.

### Incucyte assays-Proliferation and tumor sphere formation

10,000 cells were plated in a 96 well plate in triplicates. Brightfield images were collected by Incucyte every 3 hours for a total of 72 hours, until cells reached confluency. For the tumor shphere assay, 100 μL of Matrigel (Corning) was coated on the well and cured at 37C for 30 minutes. 1000 cells were embedded into the Matrigel and allowed to form tumor-spheres for ∼100 hours. Image analysis was done to quantify % confluence, counts and size of tumorspheres and plotted as a function of time.

### Efferocytosis assay

Differentiated BMDMs were seeded in a 12-well plate and cultured in IMDM medium containing 0.5% HI FBS, with or without Dexamethasone, for 18 hours. Apoptotic cells were generated by treating Jurkat cells with 1 µM Staurosporine (FUJIFILM) for 3 hours at 37°C in RPMI medium without serum. After treatment, apoptotic cells were washed and labeled with 100 ng/mL pHrodo Red (ThermoFisher) for 30 minutes, followed by two washes with PBS containing 1% BSA and 1 mM EDTA, and one wash with IMDM. The labeled apoptotic cells were then resuspended in IMDM supplemented with 10% FBS and added to the plated macrophages at a 3:1 ratio. The co-culture was incubated for 45 minutes at 37°C. After incubation, macrophages were washed twice with PBS and detached by gentle scraping. Efferocytosis was assessed by flow cytometry, measuring the percentage of pHrodo^+^ cells within the CD11b^+^ F4/80^+^ macrophage population.

### Nanostring RNA profiling

EO771 tumors were excised on day 15 after implantation and flash-frozen in liquid nitrogen. The tumors were digested in Trizol using a rotor-stator homogenizer or the Precellys tissue homogenizer. RNA was extracted using chloroform extraction. The upper aqueous phase was carefully aspirated, and the RNA was precipitated using isopropanol and incubating the mixture at −80° C for 2 hours. The RNA pellet was collected by centrifugation and was washed twice with 70% ethanol. The RNA pellet was resuspended in RNAse free H_2_O and quality and quantity determined by measuring the OD at A260, A230 and A280 using Nanodrop. The Nanostring gene expression analysis was carried out on the nCounter Max using the Pan-Cancer IO 360 panel by the Rutgers Immune Monitoring and Flow Cytometry Core. The data analysis was done using the Nanostring Rosalind and nSolver platforms.

### Statistical analysis

All in vitro experiments were repeated at-least 3 times. All differences between groups in all in vivo experiments were examined for statistical significance using a two-tailed Student t test and two-way ANOVA or Mixed effects model to compare multiple groups. GraphPad Prism software was used to perform statistical analysis and p < 0.05 was considered significant.

## Results

### PS is chronically externalized in the tumor micro-environment in E0771 orthotopically grafted tumor-bearing mice

Previous studies employing an E0771 orthotopic syngeneic breast cancer model showed therapeutic efficacy when either PS-targeting antibodies (Bavituximab) or PS receptors (Mertk) were administered in combination with anti-PD1 checkpoint therapeutics, suggesting an inhibitory PS and PS-Receptor axis that crosstalks to T cells is functional in this model ^38, 45, 46^. However, despite the importance of *inhibitory* PS signaling in the tumor microenvironment, it is still unclear (i) the mechanisms by which PS becomes chronically dysregulated in solid cancers and (ii) the cell type(s) that contribute to PS externalization and inhibitory signaling to achieve immune evasion. Here, in an attempt to investigate the most proximal cell intrinsic events that contribute functionally to PS externalization and immune signaling, we targeted two preeminent PS scramblases expressed on the E0771 cells that include the caspase-activated scramblase Xkr8 ^13–15^ and the calcium activated scramblase TMEM16F ^16, 47, 48^. To verify that the E0771 (orthotopic tumor) model displayed constitutively exposed PS, we first cloned, expressed, and purified PS-targeting antibodies 11.31, 1N11, and Bavituximab from Expi293 cells ^49^. We also employed a truncated (GLA+EGF)-Fc (mouse IgG2a) PS-binding Fc fusion domains that preferentially bind to PS (Gla-Fc-1; Fig.1.A.) ^50^. As previously noted, the purified antibodies 1N11 and Bavituximab bind to PS indirectly through β2GPI, whereas 11.31 and Gla-EGF(4) binds directly to PS ^49^.

**Figure 1:**
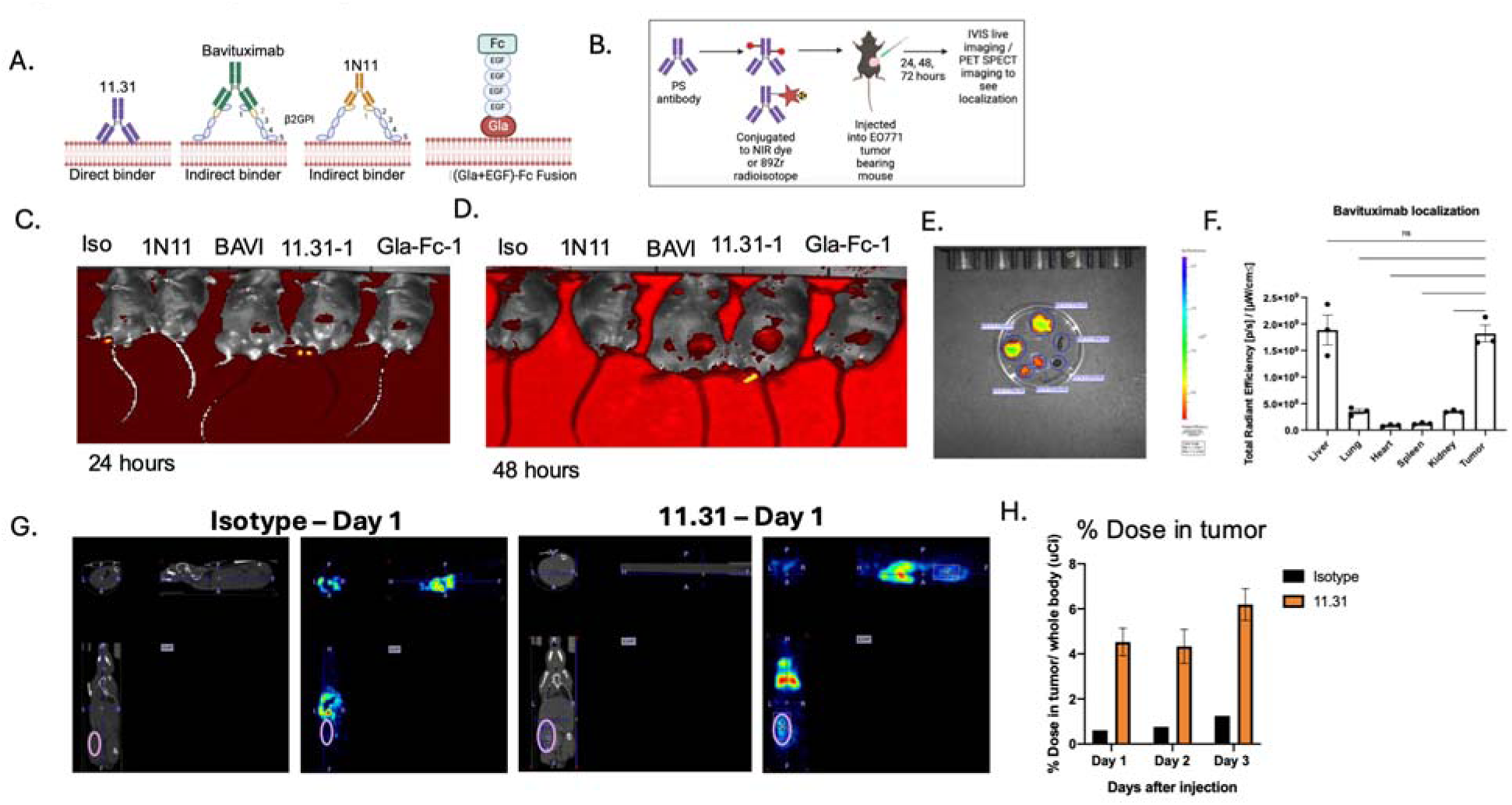
*In vivo* localization and biodistribution of PS-targeting antibodies in E0771 tumor-bearing mice. (A) Schematic showing the antibodies and recombinant proteins used in this study. PS-targeting antibodies 1N11, 11.31, and Bavituximab, as well as a truncated PS-binding modality (GLA+EGF) domain, were expressed and purified for PS binding. (B) Schematic illustrating the experimental setup for tracking PS externalization in E0771 orthotopically grafted tumor-bearing mice using PS-targeting antibodies labeled with near-infrared (NIR) dye or ^89^Zr. (C) IVIS imaging at 24 hours post-injection showing tumor localization of PS-targeting antibodies (11.31, 1N11, Bavituximab and Gla-EGF-Fc) and (D) IVIS imaging at 48 hours post-injection showing continued localization of PS-targeting antibodies in the tumor, while isotype control showed minimal signal. (E) Tissue distribution analysis of Bavituximab at 72 hours after injection, with highest localization observed in tumors and liver, followed by kidney, and minimal in lungs, spleen, and heart. (F) Quantification of Bavituximab localization in tumors compared to other tissues, indicating significant tumor-specific PS exposure (Ordinary One way ANOVA, n=3, * =p<0.05, ** =p < 0.01, *** = p <0.001, **** = p < 0.0001). (G) PET/CT imaging of ^89^Zr-labeled 11.31 antibody at 24 hours post-injection in EO771 tumor-bearing mice, with an isotype control showing lower localization. (H) Quantification of 11.31 localization showing a sustained increase in tumor localization for up to 72 hours compared to isotype control, with liver accumulation observed for both 11.31 and isotype.

To characterize *in vivo* PS externalization in the tumor microenvironment, we labelled aforementioned PS targeting antibodies with an NIR dye or radioisotope Zr-89 to track their biodistribution in tumor bearing mice. *In vivo* imaging methods including IVIS and PET/CT were employed to monitor PS antibody localization in the E0771 tumor-bearing mice (Fig.1.B). IVIS imaging showed *in vivo* localization of PS targeting antibodies and the Gla-EGF fusion protein to the tumors at 24 (Fig.1.C.) and 48 hours (Fig.1.D.) after intraperitoneal injection, while Isotype antibody showed minimal tumor localization. Tumors, liver, spleen lungs, heart and kidneys were harvested at 72 hours after injection and subjected to IVIS. Tissue distribution of Bavituximab was seen to be most prominent in tumors and liver, followed by kidney, and least in lungs, spleen and heart (Fig.1.E.). Antibody accumulation in the liver and kidneys could be due to immune complex formation. Tumors notably showed high levels of antibody localization of Bavituximab, suggesting that tumors externalize PS, and that the PS is constitutive and available for PS targeting by therapeutic modalities (Fig.1.F.).

To verify the above findings, *in vivo* imaging studies were also conducted with ^89^Zr-labelled direct PS binding antibody 11.31 and assessed using PET/CT imaging. 11.31 and isotype were labelled with ^89^Zr (chosen due to its long half-life of 78.4 hours, allowing us to track antibody localization for longer times up to 72 hours). PET/CT imaging showed localization of 11.31 to EO771 tumor tissues at 24, 48 and 72 hours after injection, compared to isotype (Fig 1.G.). As noted, accumulation in the liver was observed in both isotype and 11.31; however, the tumors (circled) showed a notable increased localization of the 11.31 antibody that was sustained for 72 hours after injection compared to the isotype (Fig.1.H.). Taken together, our results indicate that E0771 tumors harbor *robust* PS externalization in orthotopic tumor-bearing mice, as previously shown by other groups ^38^.

While PS-targeting antibodies, shown here and elsewhere, are known to hone to the tumor microenvironment in solid cancers, it is still not clear the functional contribution of tumor cells versus tumor vasculature and stroma cells towards immune evasion ^18, 51^. Indeed, while previous studies reported that PS antibodies can bind to the tumor vasculature ^32, 40, 52^, the contribution of tumor cells functioning as a cell intrinsic event to PS externalization and immune escape mechanisms in the tumor microenvironment is not defined. To investigate the cell intrinsic role of PS on tumor cells, we explored targeting PS scramblases TMEM16F and Xkr8 specifically on E0771 tumor cells by CRISPR/Cas9 gene editing.

### Ablation of Xkr8 on E0771 tumor cells blocks PS externalization on dying cells without altering cell intrinsic oncogenic properties

To investigate the expression patterns of Xkr family members including Xkr4, Xkr8, and Xkr9, (all of which contain caspase recognition sites for activation), we screened the E0771 cells by q-RT-PCR (Fig. 2A), Notably, of these isoforms the Xkr8 isoform was predominantly expressed at the RNA level (Fig. 2A), suggesting a predominant role of Xkr8 in PS scrambling and externalization. To assess biological function, Xkr8 was targeted by CRISPR/Cas9 gene editing, after which single cell clones were selected and passaged. Successful knockout was verified using a custom polyclonal rabbit anti-mouse Xkr8 antibody or by a surveyor PCR assay to interrogate the gene locus (Fig. 2B, SFig 2.A). To assess functional impairment of PS externalization during apoptosis, we treated the E0771 Xkr8 KO cells with 100 nM Staurosporine, an ATP kinase inhibitor and inducer of apoptosis, and assessed PS externalization in real time using Incucyte and fluorescently labelled Annexin-V as a function of time. As shown in Fig. 2C, Xkr8 knockout cells showed a dramatic reduction in the intensity of fluorescently labelled Annexin V compared to WT cells, indicating PS externalization can be uncoupled from caspase-mediated apoptosis in the E0771 cells. Interestingly, however, E0771 Xkr8 knockout cells did not show apparent defects in tumor cell proliferation (Fig.2.D.) or tumorsphere formation in Matrigel as shown by tumorsphere size (Fig.2.E.) and tumorsphere count (Fig.2.F.), or defects in the immediate Gas6-maediated activation of Akt (SFig 2B) suggesting the Xkr8 does not impinge on the cell intrinsic oncogenic features of these cells in cell culture.

**Figure 2:**
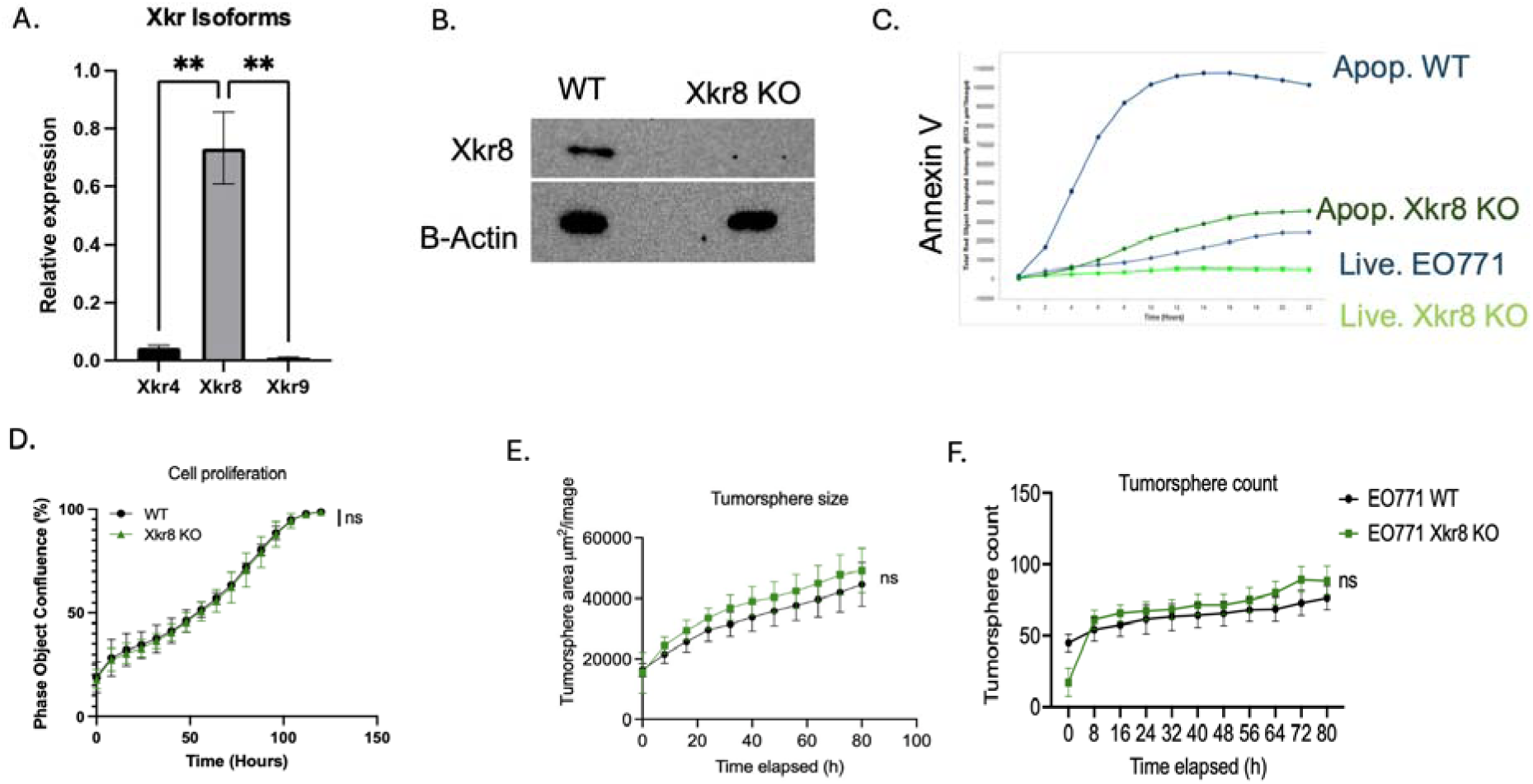
Ablation of Xkr8 on EO771 tumor cells blocks PS externalization during apoptosis without affecting intrinsic oncogenic properties. (A) q-RT-PCR analysis of Xkr isoform expression (Xkr4, Xkr8, Xkr9) in EO771 tumor cells. Xkr8 is the most highly expressed isoform among the Xkr family (Ordinary One way ANOVA, n=4, * =p<0.05, ** =p < 0.01, *** = p <0.001, **** = p < 0.0001. (B) Validation of Xkr8 knockout (KO) in EO771 tumor cells using western blotting with a custom polyclonal anti-Xkr8 antibody (Abclonal). Complete ablation of Xkr8 is observed in KO clones. (C) Real-time measurement of PS externalization in Xkr8 KO and wild-type (WT) EO771 cells upon apoptosis induction with 100 nM Staurosporine using fluorescently labelled Annexin-V, as measured by Incucyte. (D) Cell proliferation assay comparing Xkr8 KO and WT EO771 tumor cells as measured by bright field phase object confluence % by Incucyte. Tumor-sphere formation assay in Matrigel comparing Xkr8 KO and WT EO771 tumor cells was also measured by Incucyte. (E) Quantification of tumor-sphere area μM^2^ /Image and (F) tumor sphere counts were compared for Xkr8 KO and WT EO771 tumor-spheres.

### Xkr8 knockout on tumor cells reduces tumor growth *in vivo* in the orthotopic E0771 breast cancer model

To investigate *in vivo* tumor growth in an immunocompetent mouse model, we used the syngeneic orthotopic breast cancer as previously described ^45, 46^. As shown in Fig. 3, when EO771 WT cells or EO771 Xkr8 KO cells were injected into mammary gland fat pads of C57BL6 WT mice for longitudinal tumor volume studies (Fig 3.A.), the Xkr8 KO tumors exhibited significant reduction in tumor growth compared to WT tumors, as evident by tumor volume (Fig.3.B.), tumor weight (Fig.3.C.) and spleen weight (Fig.3.D.). To examine the dependence of an immune component in the tumor immune microenvironment, we compared the immunocompetent background above (Fig. 3A-D) with NSG and RAG1 immune deficient mouse models. When WT E0771 or Xkr8 KO cells were injected into NSG mice, which have a dysfunctional adaptive immune system, the KO tumors grew at a similar rate compared to the EO771 WT tumors, as shown by tumor volume (Fig.3.E.) and tumor weight (Fig.3.F.). Similar observations were noted in the Rag1 KO mice, which do not have matured T and B cells, whereby the Xkr8 KO tumors grew similarly as WT evident by both tumor volume (Fig. 3.G) and tumor weight (Fig.3.H.). Together, these observations suggest an active role of an immune component in the tumor regression in Xkr8 KO tumors.

**Figure 3:**
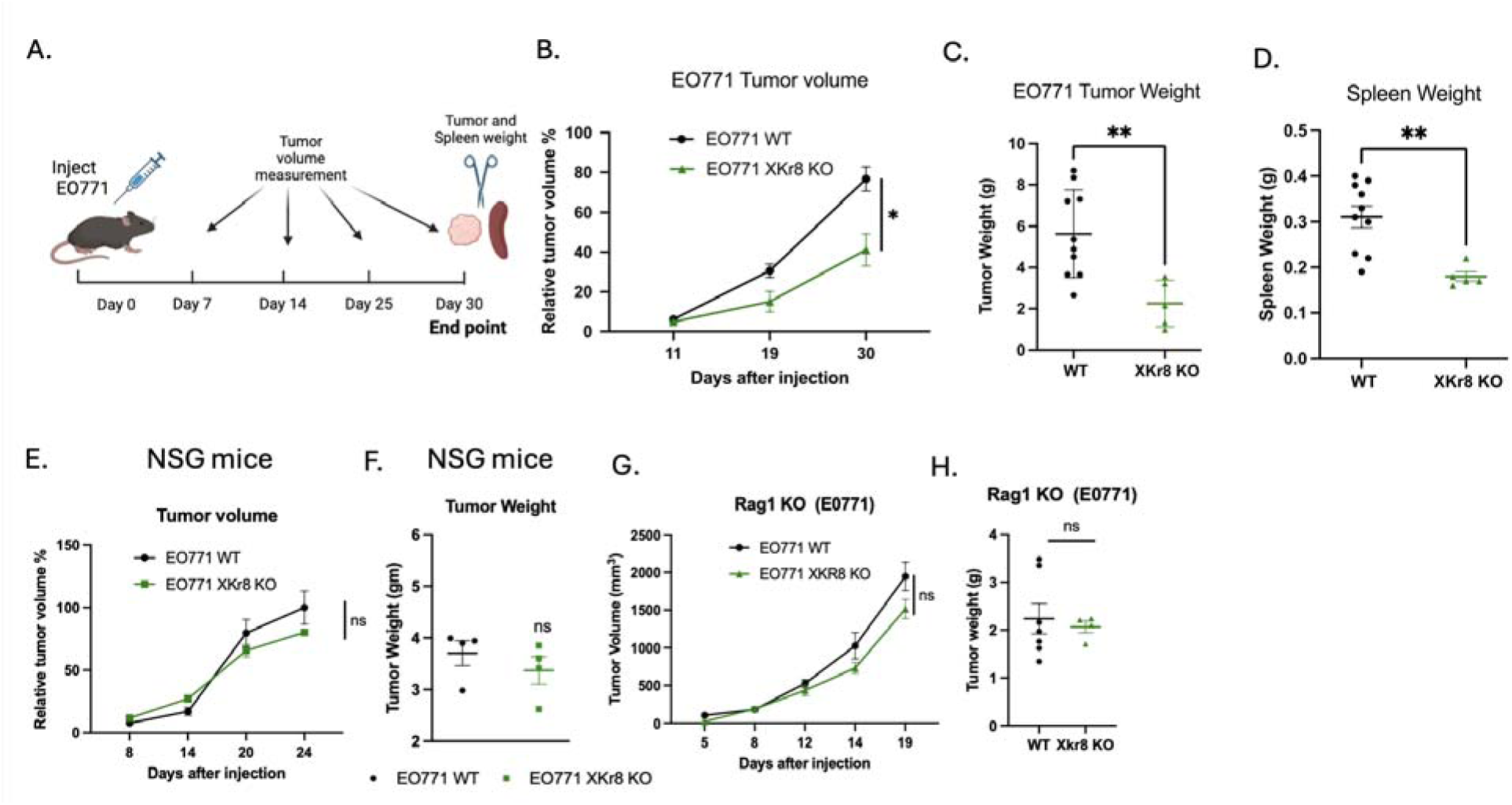
Xkr8 knockout reduces tumor growth in an orthotopic model of breast cancer. (A) Experimental timeline for orthotopic injection of EO771 wild-type (WT) or Xkr8 knockout (KO) tumor cells into the mammary fat pads of C57BL/6 wild-type (WT) mice. Tumor volumes were measured periodically to assess growth. Tumor weights and spleen weights were measured at end. (B) Tumor volumes of E0771 WT and Xkr8 KO tumors at various time points point (Mean values ± SD, Two-way ANOVA, Mixed effects model, n=15 * =p<0.05, ** =p < 0.01, *** = p <0.001, **** = p < 0.0001). (C) Tumor weights of EO771 WT and Xkr8 KO tumors at the time of sacrifice point (Mean values ± SD, One-way ANOVA, n=15 * =p<0.05, ** =p < 0.01, *** = p <0.001, **** = p < 0.0001). (D) Spleen weights of EO771 WT and Xkr8 KO tumor-bearing mice point (Mean values ± SD, One-way ANOVA, n=15 * =p<0.05, ** =p < 0.01, *** = p <0.001, **** = p < 0.0001). (E) Tumor volumes in immune-deficient NSG mice, which have a dysfunctional adaptive immune system, injected with EO771 WT or Xkr8 KO cells (Mean values ± SD, Two-way ANOVA, Mixed effects model, n=9 * =p<0.05, ** =p < 0.01, *** = p <0.001, **** = p < 0.0001). (F) Tumor weights in NSG mice injected with EO771 WT or Xkr8 KO cells (Mean values ± SD, One-way ANOVA, n=9 * =p<0.05, ** =p < 0.01, *** = p <0.001, **** = p < 0.0001). (G) Tumor volumes (Mean values ± SD, Two-way ANOVA, Mixed effects model, n=10 * =p<0.05, ** =p < 0.01, *** = p <0.001, **** = p < 0.0001) and (H) tumor weights (Mean values ± SD, One-way ANOVA, n=9 * =p<0.05, ** =p < 0.01, *** = p <0.001, **** = p < 0.0001) in Rag1 knockout (KO) mice, which lack mature T and B cells.

### Ablation of TMEM16F on E0771 tumor cells blocks PS externalization on calcium-stressed cells without altering cell intrinsic oncogenic properties in E0771 cells

Having established Xkr8 knockout cells in vitro and *in vivo* E0771 model, we next explored the effects of calcium activated scramblases in a side-by-side comparison to Xkr8. Among the TMEM16 isoforms (16C, 16D, 16F, 16G and 16J), TMEM16F had robust expressed at the mRNA level in EO771 cells (Fig 4.A.) and therefore assessed for functional outcomes. Towards this goal, and analogous to the Xkr8 strategy, we depleted TMEM16F in EO771 cells using CRISPR-Cas9 gene editing and sorted single cell clones. Validation of the TMEM16F KOs was assessed using Western blot with an antibody specific for TMEM16F (Antibody courtesy Lily Jan, UCSF) (Fig.4.B.). To examine effects of TMEM16F on calcium-mediated PS externalization. WT and TMEM16F knockout E0771 cells were treated with the calcium ionophore A23187, after which PS externalization in response to calcium influx was monitored using Annexin V-FITC binding by microscopy (Fig. 4.C.) or flow cytometry (Fig.4.D.). Notably, while the EO771 WT cells externalized PS within 20 minutes, by contrast the TMEM16F KO cells showed a dramatic reduction and marginalized PS externalization under these conditions (Fig 4.C,D.). Analogous to the assessment of Xkr8 in cell intrinsic oncogenic features, TMEM16F knockout cells also did not show defects in tumor cell proliferation (Fig.4.E.) or tumor-sphere formation in Matrigel, as shown by tumor-sphere size (Fig. 4.F.) and tumor-sphere count (Fig 4.G.), or defects in the immediate Gas6-maediated activation of Akt (SFig 2B).

**Figure 4:**
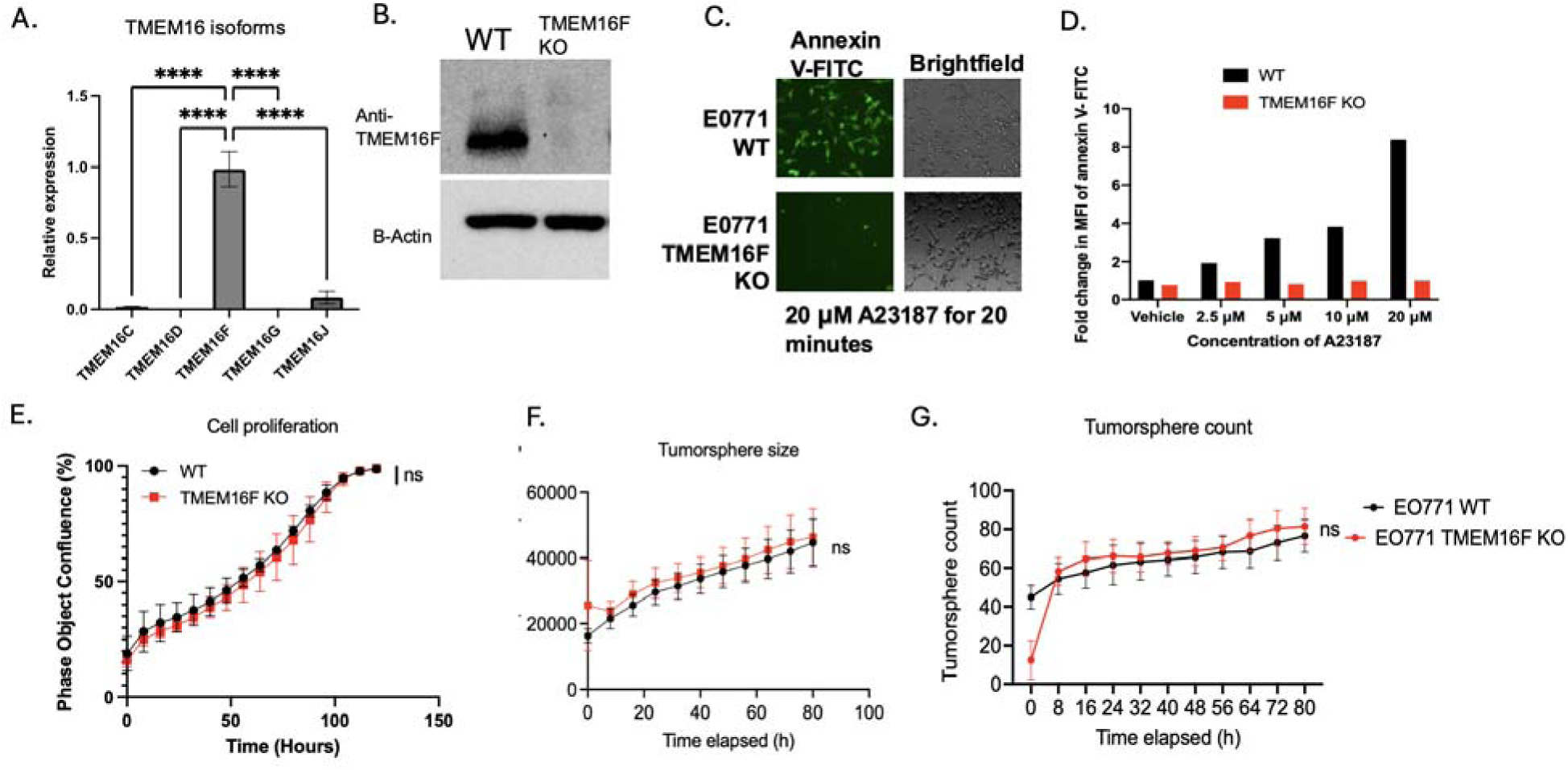
TMEM16F knockout prevents PS externalization in response to calcium stress without altering intrinsic oncogenic properties. (A) q-RT-PCR analysis showing the relative expression of TMEM16 isoforms (16C, 16D, 16F, 16G, and 16J) in E0771 cells (Mean Relative gene expression values ± SD, Ordinary One-way ANOVA, n=3, * =p<0.05, ** =p < 0.01, *** = p <0.001, **** = p < 0.0001). (B) Western blot validation of TMEM16F knockout (KO) in EO771 cells. The absence of TMEM16F is confirmed in the KO clones using a custom antibody (courtesy of Lily Jan, UCSF). (C) Fluorescent microscopic analysis of PS externalization in response to calcium influx using A23187 (calcium ionophore) and Annexin V-FITC staining. E0771 wild-type (WT) cells show robust PS externalization, while TMEM16F KO cells exhibit significantly reduced PS externalization. (D) Fold change in MFI for Annexin V-PE staining by flow cytometry of PS externalization in EO771 WT and TMEM16F KO cells upon treatment with A23187. (E) Cell proliferation assay comparing EO771 WT and TMEM16F KO cells using Incucyte as measured by bright field phase object confluence % by Incucyte. Tumor-sphere formation assay in Matrigel comparing TMEM16F KO and WT EO771 tumor cells was also measured by Incucyte. (F) Quantification of tumor-sphere area μM^2^ /Image and (G) tumor sphere counts were compared for TMEM16F KO and WT EO771 tumor-spheres.

### TMEM16F KO on tumor cells reduce tumor growth in the E0771 orthotopic model of breast cancer

To assess the effects of TMEM16F knockout on tumor growth and the tumor microenvironment and compare effects to that of the Xkr8 knockout model described above (Fig. 3), EO771 WT cells or EO771 TMEM16F KO cells were transplanted orthotopically into the mammary gland fat pads of C57BL/6 WT mice. Interestingly, in a similar capacity to Xkr8, the TMEM16F KO tumors also showed significant reduction in tumor growth compared to WT tumors (employing 2 independent clones for validation), as evident by the decrease in tumor volume (Fig.5.A.D.), tumor weight (Fig.5.B.,E.) and spleen weight (Fig.5.C.), as well as increased overall survival of mice with TMEM16F KO tumors, as indicated by a Kaplan Meier curve (Fig 5.F.) Similar results were obtained using an MC38 flank colon cancer model, whereby TMEM16F knockout also reduced flank tumor growth, as observed by tumor volume (Fig.5.G.), tumor weight (Fig 5.H.) and spleen weight (Fig 5.I.). These data demonstrate that TMEM16F affects tumor progression in both orthotopic and flank models. However, TMEM16F KO and WT E0771 tumor growth were comparable in immunocompromised NSG mice, as evident by no significant reduction in the tumor volume (Fig 5.J.) and tumor weight (Fig 5.K.). Interesting, unlike the Xkr8 knockout phenotype, the TMEM16F KO tumors showed a slight reduction in tumor growth in Rag KO mice (which lack mature T and B lymphocytes), with modest effects in tumor volume (Fig 5.L.) and tumor weight (Fig 5.M.), possibly suggesting that tumor reduction due to Xkr8 KO is more dependent on the mature T and B cells, whereas, in the TMEM16F KO tumors, there is a combined effect of the immune system and other factors, such as angiogenesis and modulation of the extracellular matrix, driving the reduction in tumor progression.

**Figure 5:**
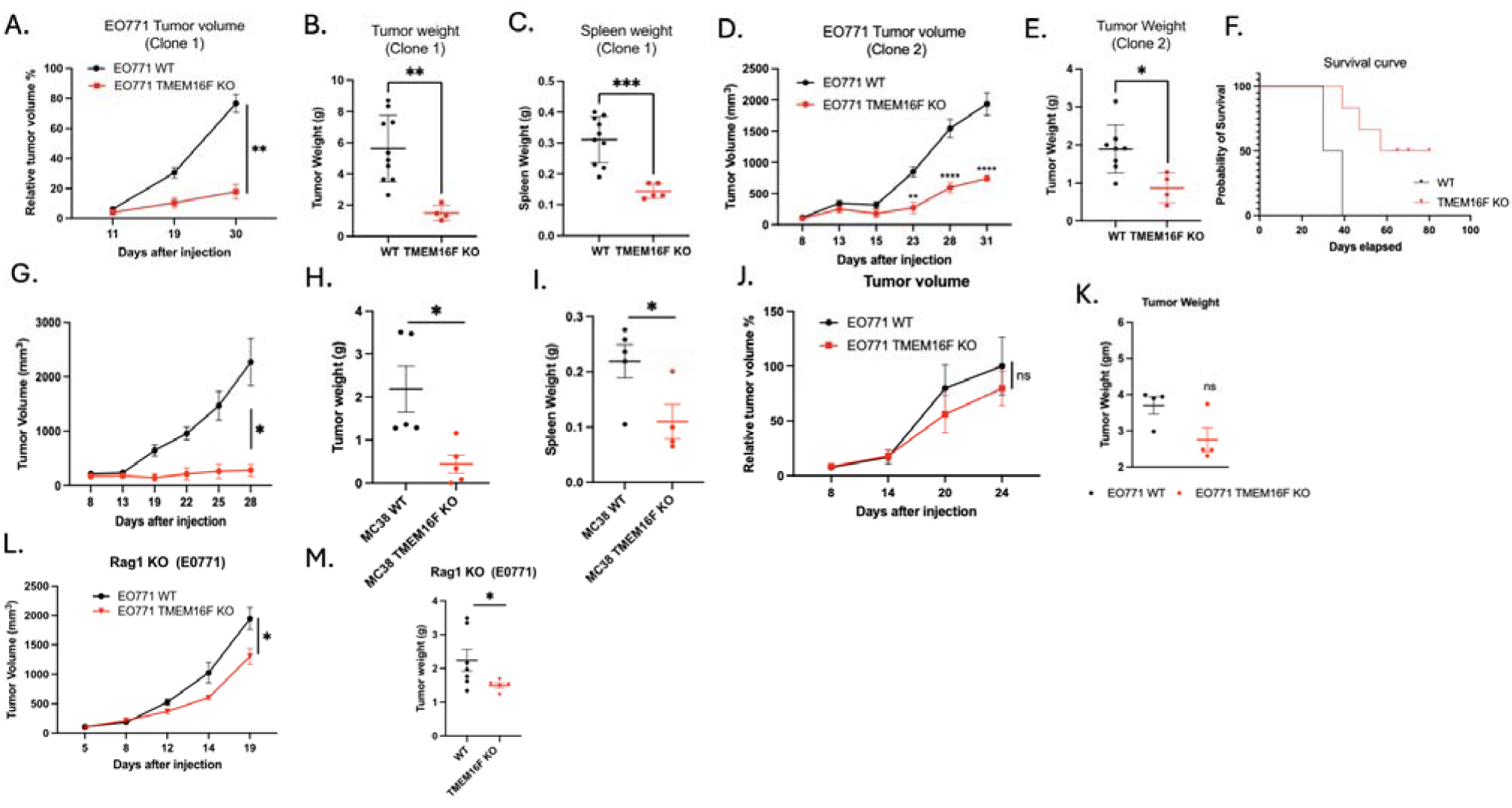
TMEM16F knockout reduces tumor growth in syngeneic orthotopic models of breast and colon cancer. (A) Tumor volume (mm^3^) measurements from C57BL6 WT mice injected with E0771 WT or TMEM16F KO cells into mammary fat pads (Mean values ± SD, Two-way ANOVA, Mixed effects model, n=15 * =p<0.05, ** =p 0.01, *** = p <0.001, **** = p < 0.0001). (B) Tumor weight (g) measurements from the same C57BL6 WT mice showing reduced tumor weight in TMEM16F KO tumors compared to WT tumors (Mean values ± SD, One-way ANOVA, n=15 * =p<0.05, ** =p < 0.01, *** = p <0.001, **** = p < 0.0001).(C) Spleen weight (g) measurements from C57BL6 WT mice injected with EO771 WT or TMEM16F KO cells (Mean values ± SD, One-way ANOVA, n=15 * =p<0.05, ** =p < 0.01, *** = p <0.001, **** = p < 0.0001). (D) Tumor volume (mm^3^)(Mean values ± SD, Two-way ANOVA, Mixed effects model, n=10 * =p<0.05, ** =p < 0.01, *** = p <0.001, **** = p < 0.0001), (E) tumor weight (g) (Mean values ± SD, One-way ANOVA, n=10 * =p<0.05, ** =p < 0.01, *** = p <0.001, **** = p < 0.0001) and (F) spleen weight (g) (Mean values ± SD, One-way ANOVA, n=10 * =p<0.05, ** =p < 0.01, *** = p <0.001, **** = p < 0.0001) measurements from an independent clone: 2 of E0771 TMEM16F KO cells, showing consistent reduction in tumor growth compared to WT tumors. (F) Kaplan-Meier survival curve showing improved survival in mice bearing EO77 TMEM16F KO tumors compared to those with WT tumors, indicating the beneficial effect of TMEM16F knockout on survival (n=10). (G) Tumor volume, (Mean values ± SD, Two-way ANOVA, Mixed effects model, n=10 * =p<0.05, ** =p < 0.01, *** = p <0.001, **** = p < 0.0001) (H) tumor weight (Mean values ± SD, One-way ANOVA, n=10 * =p<0.05, ** =p < 0.01, *** = p <0.001, **** = p < 0.0001), and (I) spleen weight measurements from MC38 colon cancer model in C57BL6 WT mice (Mean values ± SD, One-way ANOVA, n=15 * =p<0.05, ** =p < 0.01, *** = p <0.001, **** = p < 0.0001). (J) Tumor volume (Mean values ± SD, Two-way ANOVA, n=9 * =p<0.05, ** =p < 0.01, *** = p <0.001, **** = p < 0.0001)and (K) tumor weight (g) (Mean values ± SD, One-way ANOVA, n=9 * =p<0.05, ** =p < 0.01, *** = p <0.001, **** = p < 0.0001) measurements in NSG mice injected with EO771 WT or TMEM16F KO cells, showing no significant difference between the two groups. (L) Tumor volume (Mean values ± SD, Two-way ANOVA, Mixed effects model, n=10 * =p<0.05, ** =p < 0.01, *** = p <0.001, **** = p < 0.0001)and (M) tumor weight measurements in Rag1 KO mice injected with E0771 WT or TMEM16F KO cells (Mean values ± SD, One-way ANOVA, n=15 * =p<0.05, ** =p < 0.01, *** = p <0.001, **** = p < 0.0001).

### Distinct phenotypic outcomes in the Xkr8 and TMEM16F knockout E0771 cells

The pioneering studies by Nagata and colleagues described the distinct biochemical and molecular features of Xkr8 and TMEM16F as caspase and calcium-activated scramblases respectively. This interpretation is supported in the E0771 model employed here, showing that the E0771 cells lacking Xkr8 showed reduction in externalizing PS during staurosporine (caspase)-mediated apoptosis, while the E0771 cells lacking TMEM16F expression showed defective PS externalization during ionophore (calcium)-induced cytosolic calcium elevation.

To access the mechanistic and functional consequences of the scramblase knockouts, we examined *in vitro* efferocytosis using pHrodo-labeled apoptotic E0771 WT, Xkr8 KO and TMEM16F KO cells when co-cultured with bone-marrow derived macrophages (BMDMs) (Fig. 6A). As indicated in Fig. 6B, whereas the BMDMs efficiently engulfed apoptotic WT and apoptotic TMEM16F KO cells, efferocytosis was notedly reduced when Xkr8 KO apoptotic cells were co-cultured with macrophages. Moreover, and analogous to the observations in Fig. 2C, when assessed by flow cytometry for Annexin V-FITC, only the Xkr8 KO cells, but not the TMEM16F KO cells, showed suppression in apoptosis-mediated PS externalization (Fig. 6C-D). TMEM16F KO apoptotic cells were efficiently engulfed by BMDMs, suggesting distinct functions for scramblases, showing that Xkr8 mediated PS externalization on apoptotic tumor cells is essential for recognition by BMDMs (Fig.6C). To corroborate this, we also show that PS is externalized by WT and TMEM16F KO apoptotic cells and not by Xkr8 KO apoptotic cells (Fig. 6C, 6D).

**Figure 6.**
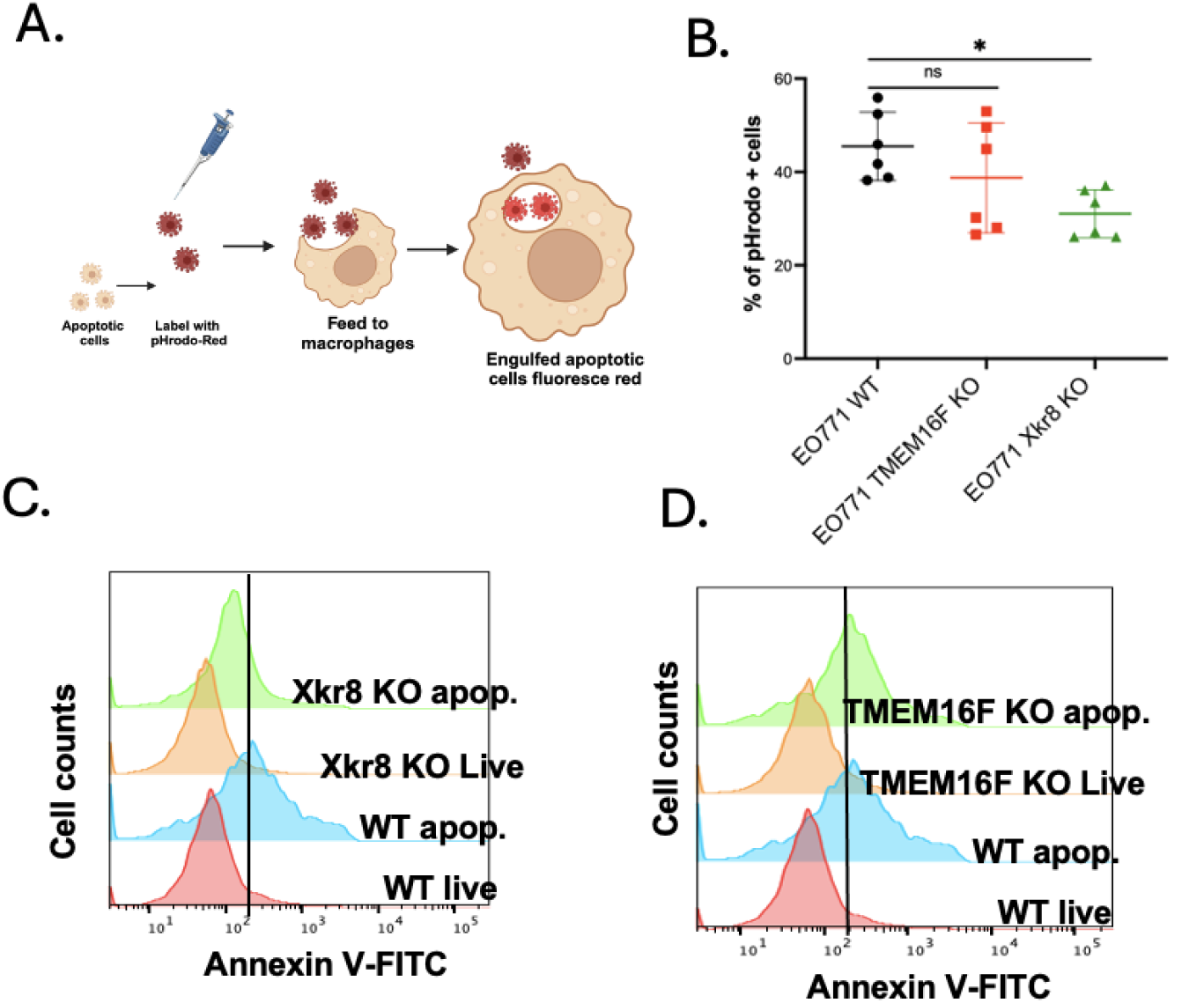
Apoptotic tumor cells induce efferocytosis by Xkr8. (A) Schematic showing the workflow for efferocytosis assays, wherein apoptotic E0771 cells (WT, TMEM16F KO and Xkr8 KO) were labelled with pHrodo red, fed to bone marrow derived macrophages, and after 3 hours, % of uptake was measured by flow cytometry. (B) Rate of efferocytosis as shown by % of pHrodo red + cells (Mean ± SD, One-way ANOVA, n=6 * =p<0.05, ** =p < 0.01, *** = p <0.001, **** = p < 0.0001). (C) PS externalization was disrupted in Xkr8 KO apoptotic cells, but not in WT and TMEM16F KO apoptotic cells (D), providing evidence that Xkr8-mediated PS externalization is essential for efficient recognition and phagocytosis by BMDMs.

The findings that TMEM16F KO still retained their capacity to undergo caspase-mediated cell death and PS externalization suggest TMEM16F functions at a distinct level that impinges on calcium-mediated PS externalization. To interrogate the relationship between elevated intracellular calcium and calcium-mediated PS externalization, we generated a calcium reporter cell line was made by stably transfecting WT or TMEM16F knockout EO771 cells plasmid expressing a genetically encoded calcium indicator, GCaMP6f, targeted to the cytosol (Fig 7.J.). The GCaMP family of calcium indicators were developed from circularly permutated green fluorescent protein (cpGFP) linked to an engineered calmodulin (CaM) and a CaM-binding M13 peptide. The binding of calcium to calmodulin (Kd ≈ 380 nm) triggers a conformational change within the cpGFP chromophore resulting in a large increase in fluorescence intensity at 485-nm excitation. ^53^. As noted in Fig. 7A, while intracellular calcium is equally upregulated in the WT and TMEM16F KO cells upon treatment with calcium ionophore, the PS externalization was severely impaired in the TMEM16F KO cells (Fig. 7.B), but not the WT cells, indicating that PS can be functionally dissociated from the elevated intracellular calcium by ablating TMEM16F.

**Figure 7.**
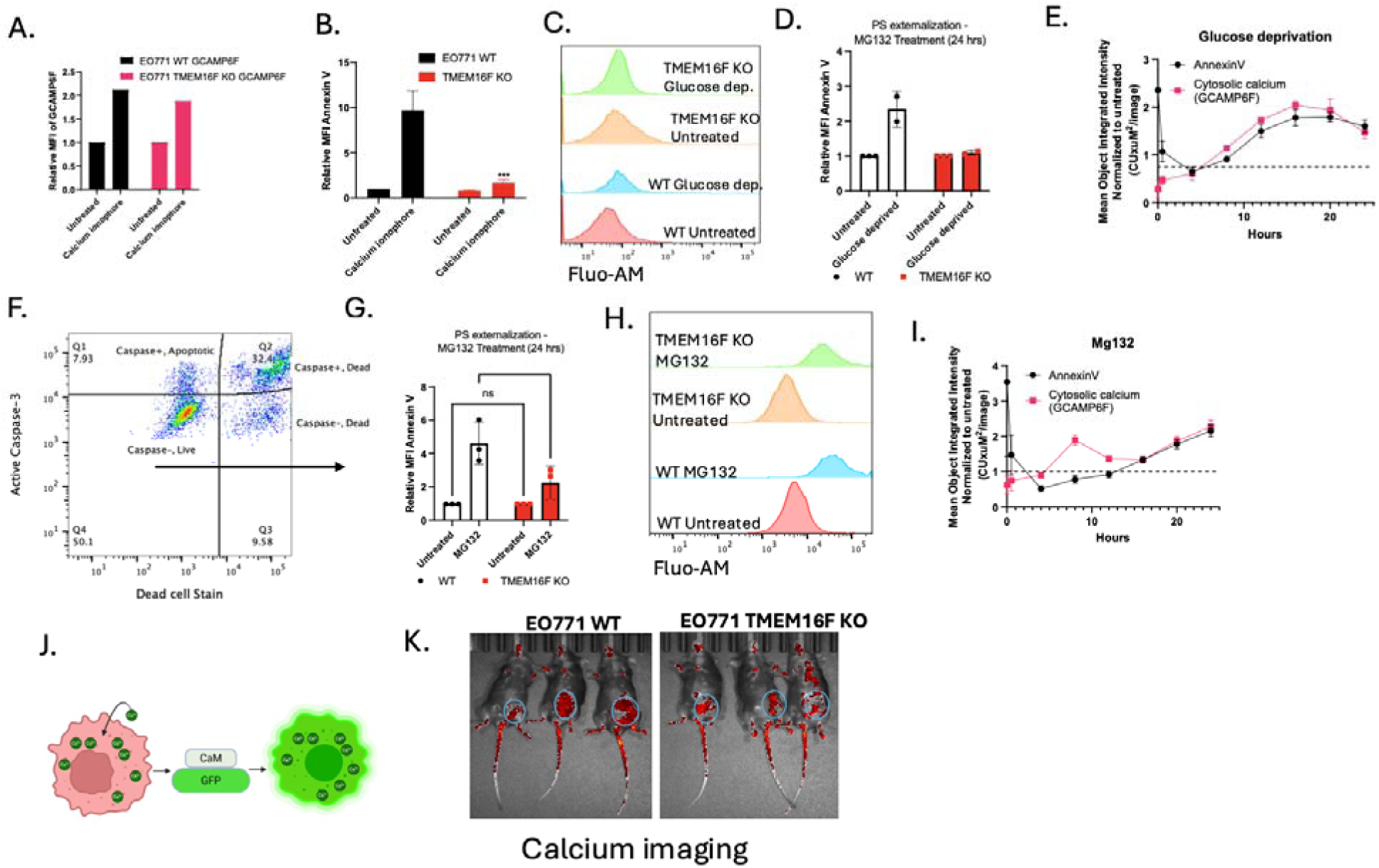
Intracellular calcium regulation and PS externalization in tumor cells. (A) Intracellular calcium levels were upregulated in both WT and TMEM16F KO EO771 cells upon treatment with calcium ionophore, as indicated by the increased calmodulin-GFP fluorescence. (B) PS externalization as measured by relative MFI of Annexin V staining in calcium ionophore treated WT and TMEM16F KO cells (Mean values ± SD, One-way ANOVA, n=3 * =p<0.05, ** =p < 0.01, *** = p <0.001, **** = p < 0.0001). (C) Flow plots showing intracellular calcium levels as shown by Fluo-AM staining. (D) PS externalization as measured by relative MFI of Annexin V staining in WT and TMEM16F KO cells exposed to glucose deprivation for 24 hours (n=2). (E) Live cell imaging with Incucyte show calcium upregulation and PS externalization (Annexin V) for up to 24 hours with glucose deprivation. (F) Gating strategy shows that PS externalization was measured on the live cells that stained negative for active caspase 3 as well as the viability stain. (G) PS externalization on cells treated with low dose MG132, a proteasome inhibitor, as shown by relative MFI of annexin V staining (Mean values ± SD, One-way ANOVA, n=3 * =p<0.05, ** =p < 0.01, *** = p <0.001, **** = p < 0.0001). (H) Flow plots showing intracellular calcium levels upon MG132 staining as shown by Fluo-AM staining. (I) Live cell imaging with Incucyte showed calcium upregulation and PS externalization (Annexin V) for up to 24 hours MG132 treatment. (J) Schematic of the calcium reporter cell line generated by transfecting EO771 cells with a calmodulin-GFP expressing plasmid to monitor intracellular calcium levels. (K) These cells were implanted into C57BL6/WT mice and IVIS imaging for GFP expression shows the tumors harboring high intracellular calcium (circled).

Subsequently, to phenocopy effects of oncogenic endoplasmic (ER) stress, a known inducer of dysregulated and elevated cytosolic calcium, we employed two activators of ER stress that included the treatment of cells with a low dose of proteasomal inhibitor MG132, or the exposure to cells to conditions of glucose deprivation, that result in altered protein folding and protein glycosylation in the ER, respectively ^54–59^. As shown in panels Fig. 7C-E (glucose deprivation), these conditions led to increased cellular calcium and PS externalization on live (caspase negative, 7AAD negative gated cells (Fig. 7F). Similarly, treatment with low dose MG132, a proteasome inhibitor increased intracellular calcium levels in live cells (Fig’s 7G-I), also induced PS externalization via TMEM16F (Fig.7.G-I). Live cell imaging with Incucyte reveals that glucose deprivation and MG132 treatments induce sustained cytosolic calcium increases and PS externalization for up to 24 hours (Fig.7.H,I). Notably, in panels 7E and 7I, a protracted time course (10-20 hrs) showed calcium elevation and PS externalization were phenocopied (see SFig 2D, E). Moreover, when these cells were orthotopically injected into mice, elevated calcium was observed associated with the tumor mass (Fig. 7J, 7K). Taken together, these studies suggest that knockout of TMEM16F suppressed ER stress -mediated calcium dysregulation in conditions aimed to mimic oncogenic stress.

### Effects of scramblase KO on the tumor microenvironment

The tumor studies in the immune deficient mice suggested a role for the tumor immune microenvironment in the anti-tumor effect of the scramblase KO tumors. To characterize this further, we used Nanostring Gene Expression Analysis to assess differentially regulated genes in the TME. We isolated RNA from tumors grown in C57BL6 WT mice at day 15 and assessed gene expression using the mouse Pan Cancer IO 360 panel (Fig.8.A). Fig shows differentially expressed genes in Xkr8 KO and TMEM16F KO tumors compared to EO771 WT tumors respectively (Fig.8.B.). Pathway analysis determined that certain oncogenic pathways, such as angiogenesis, cell proliferation, matrix remodeling and metastasis, Notch signaling, immune cell adhesion and migration are commonly downregulated in both TMEM16F and Xkr8 KO tumors, compared to WT tumors. Upon dissecting individual gene expression, we observed TMEM16F KO and Xkr8 KO differentially regulating unique set of genes. In Xkr8 KO tumors, we observed downregulation of IL-10 (Fig.8.C.), a classical cytokine released after efferocytosis. CTLA-4, an immune checkpoint, was also seen to be downregulated in Xkr8 KO tumors, pointing to reduced immune-suppression in these tumors (Fig.8.D.). However, HMGB1, a classical DAMP that is secreted when cells undergo immunogenic cell death or secondary necrosis was observed to be upregulated in Xkr8 KO tumors (Fig.8.E.). The reduced IL-10 and increased HMGB1 in Xkr8 KO tumors indicate possibly reduced efferocytosis, and that un-cleared apoptotic cells increase inflammation.

**Figure 8:**
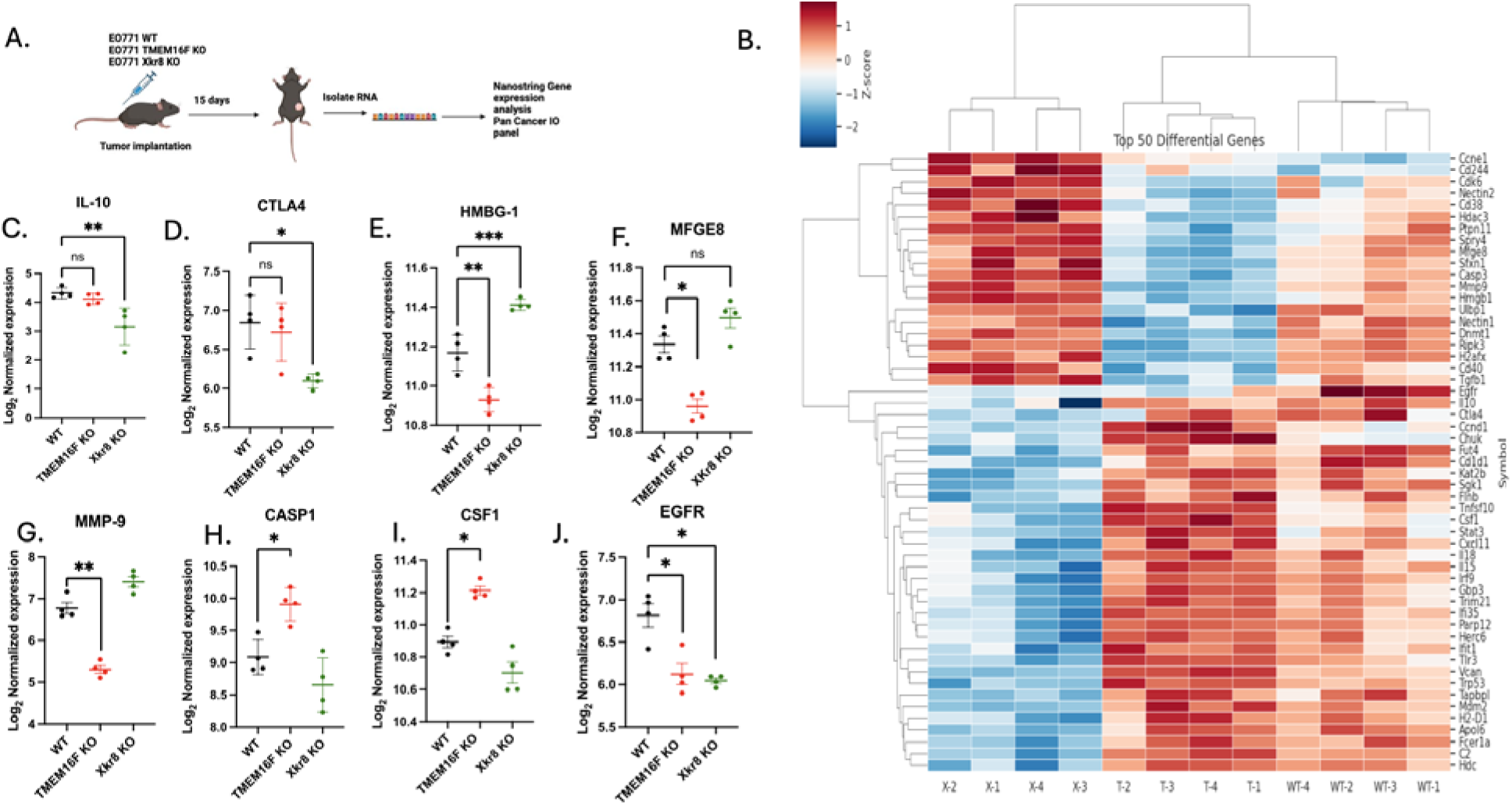
TMEM16F and XKr8 KO affect the TME through distinct pathways. (A) Schematic of the Nanostring Gene Expression Analysis used to assess the tumor immune microenvironment (TME) in tumors grown in C57BL6 WT mice at day 15 after injection. (B) Clustered heat map showing top 50 differentially regulated genes in Xkr8 KO and TMEM16F KO tumors compared to EO771 WT tumors are shown, (n=12, genes for p values<0.05 are shown and were calculated using ANOVA). WT-1,2,3,4, T-1,2,3,4 and X-1,2,3,4 correspond to EO771 WT, TMEM16F KO and Xkr8 KO tumors respectively. Gene expression analysis showed downregulation of IL-10 (C), a cytokine released after efferocytosis, and CTLA-4 (D), an immune checkpoint involved in immune suppression in Xkr8 KO tumors. In contrast, HMGB1, a damage-associated molecular pattern (DAMP) released during immunogenic cell death, was upregulated in Xkr8 KO tumors (E). (F-I) TMEM16F KO tumors exhibited downregulation of MFGE8 (F), a protein involved in PS binding and efferocytosis, and MMP9 (G), which is involved in matrix remodeling and cancer progression. Caspase-1 (H) and CSF-1 (I) were upregulated in TMEM16F KO tumors, indicating increased cell death and macrophage infiltration. (J) Both Xkr8 KO and TMEM16F KO tumors showed downregulation of EGFR expression, suggesting an impact on EGFR-expressing cell populations (Mean log_2_ normalized expression values ± SD, Two-way ANOVA, Mixed effects model, n=12 * =p<0.05, ** =p < 0.01, *** = p <0.001, **** = p < 0.0001).

Comparing EO771 WT and TMEM16F KO tumors revealed a unique differential gene expression, different from that of Xkr8 KO tumors. TMEM16F KO tumors significantly reduced MFGE8 expression (Fig.8.F.), which binds PS binding protein by its C2 domain and is a ligand for α_v_β_3_ and α_v_β_5_ integrins. TMEM16F KO tumors also showed a reduction in MMP9 (Fig.8.G.), which exacerbates cancer progression by degrading extracellular matrix proteins. Potentially consistent with this idea, the Xkr8 knockout cells had less metastasis in the NSG mice (SFig 2C). Caspase-1 and CSF-1 was also seen to be upregulated in TMEM16F KO tumors, indicating in increased cell death and macrophage infiltration respectively (Fig.8.H,I). EGFR was the common gene downregulated in both scramblase KOs, indicating that reducing PS externalization was affecting EGFR expressing cell populations (Fig.8.J.). Notably, no differences in immune cell infiltration were noted at this time point, perhaps due to lack of PS externalization in the EO771 tumors at this early time point. Taken together, while both scramblases affected tumor growth in immunocompetent mice, the mechanisms by which Xkr8 and TMEM16F impinge on immune evasion are likely distinct (Fig. 9.).

**Figure 9:**
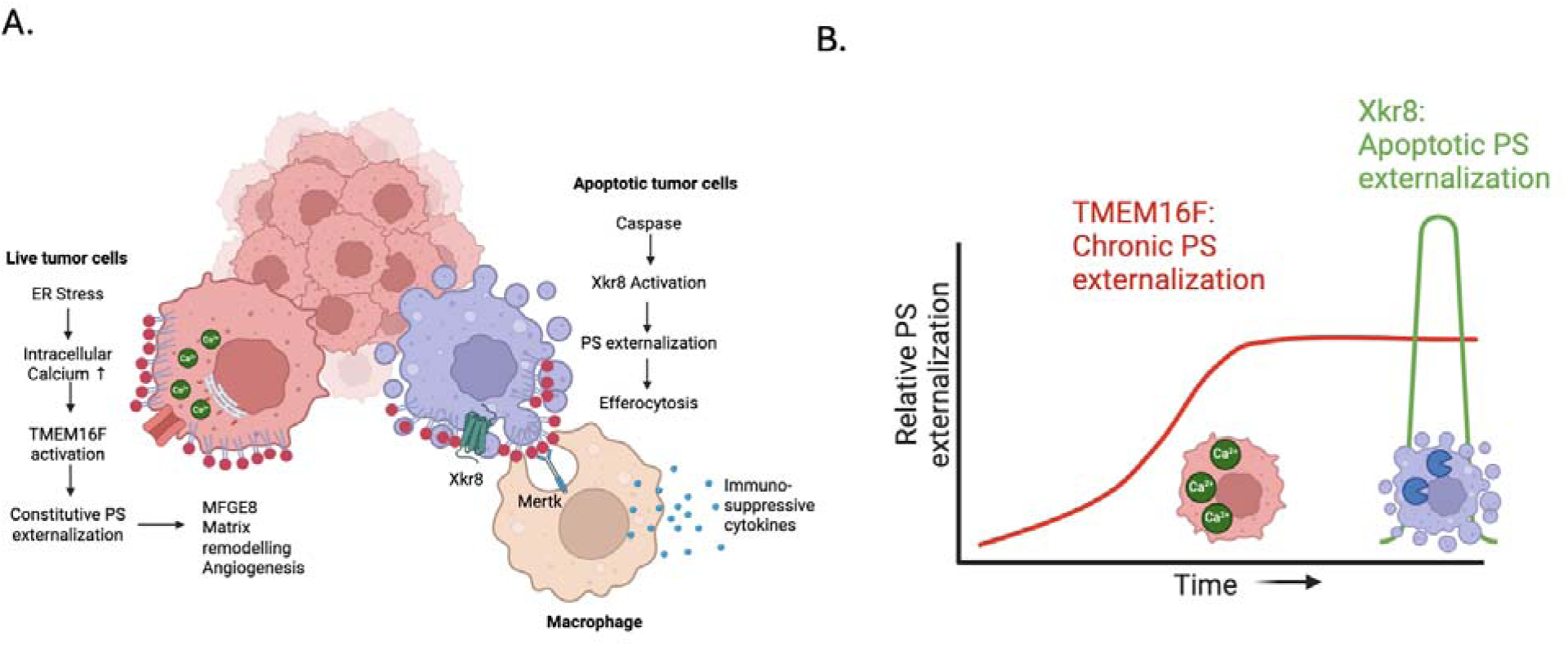
PS externalization on tumor cells by scramblases Xkr8 and TMEM16F mediate pro-tumorigenic effects by distinct pathways. (A) The working model shows that PS externalization by scramblases Xkr8 and TMEM16F mediate pro-tumorigenic effects by unique pathways, wherein Xkr8 upregulates PS exposure on apoptotic cells, leading to increased efferocytosis and secretion of immune-suppressive cytokines, whereas TMEM16F is regulated by calcium stress in live cells leading to MFGE8 mediated matrix remodeling. (B)Temporal activation of the scramblases can be explained as such: TMEM16F is activated chronically in live cells undergoing chronic ER stress and upregulated intracellular calcium, whereas Xkr8 activation is a late signal induced by caspase mediated cell death.

## Discussion

Dysregulated PS externalization has been documented in a wide array of solid tumor types as evident by both the honing and of PS-targeting monoclonal antibodies, beta-bodies, Annexins, and cationic scaffolds/peptoids to the tumor microenvironments ^18, 42, 51^ ^37^, as well as functional studies showing therapeutic utility of PS-targeting antibodies ^40, 41, 52^ ^60^ and anti-PS-Receptors (Mertk) ^45, 61^ in immune oncology, often acting in synergy with checkpoint inhibitors such as anti-PD-1. However, despite the promise of targeting PS as a universal modality in the immune-oncology of solid cancers, neither the mechanisms of PS externalization in the complex tumor microenvironment nor the repertoire of cell types that mediate persistent PS externalization are currently well articulated. In the current study, we employed the E0771 syngeneic tumor model that has been previously shown responsive to both PS-targeting Abs (Bavituximab) ^38^as well as anti-Mertk (a PS receptor of belonging to the TAM family) ^45^. We show that E0771 tumor-bearing mice with palpable tumors are targeted by two distinct PS-targeting modalities (Bavituximab and 11.31), as well as a Gla-containing PS-targeting protein, indicating the PS constitutively externalized during the tumor growth. Furthermore, E0771 cells express the two main PS scramblases isoforms Xkr8 (caspase-activated) and TMEM16F (stress and calcium activated), permitting molecular dissociation of the cancer cell intrinsic events, side-by-side, associated with apoptosis/efferocytosis (Xkr8) from the events associated with calcium dysregulation and PS externalization on oncogenic viable cells (TMEM16F). Our results provide evidence that both Xkr8 and TMEM16F knockout tumor cells suppress *in vivo* tumor growth in immunocompetent mice but in NOD/SCID or RAG1 immuno-incompetent mice. Our data support a complex and duality mechanism for PS externalization in the tumor microenvironment whereby apoptosis/efferocytosis of growing tumors and calcium-stressed PS-out viable tumor cells both likely contribute to PS-mediated immune escape and tumor progression.

Although the biochemical mechanisms by which loss of Xkr8 and TMEM16F impinge on tumor growth are still not clear, our results are consistent with a recent study by Chen and colleagues showing that co-delivery of an Xkr8 small interference RNA (siRNA) with platinum chemotherapy (to enhance apoptosis) showed increased therapeutic efficacy in an orthotopic pancreatic tumor model with an increase of proliferative NK cells and activated macrophages infiltration in the tumor microenvironment ^62^. Moreover, using single cell RNAseq, these authors showed increased infiltration of CD8+ cytotoxic T cells and reduced exhausted T cells suggesting that the loss of Xkr8 promoted a type of immunogenic death response perhaps involving the cross-presentation of tumor antigens by APCs. For example, Xkr8 deficient tumor cells showed enhanced secretion of cyclic GAMP to activate STING on neighboring cells. In the studies reported here, the Xkr8 was only deleted on the tumor cells, not other cells in the tumor microenvironment, nor did we use external chemotherapeutics to enhance tumor cell death in the *in vivo* studies. As such, we propose a model in which a finite number of proliferating tumor cells die by apoptosis in the tumor microenvironment are actively engulfed by the tumor associated macrophages and possibly other bystander phagocytes. Indeed, consistent with this idea, the E0771 cells with Xkr8 KO fail to externalize PS in vitro when treated with staurosporine, and these cells display impaired *in vitro* efferocytosis when co-cultured with bone marrow derived macrophages. Additionally, the Nanostring gene expression profiling revealed an efferocytic signature including decreased PS opsonin molecule MFG-E8, which has been shown to link apoptotic cells to PS-mediated efferocytosis with integrins ^63–65^. Interestingly, since recent studies suggest that chemotherapies can induce Xkr8 at both the transcriptional and translation levels, further studies are needed to assess the synergy of Xkr8 targeting modalities with chemotherapies or inducers of immunogenic death.

While the increased immunogenic outcomes associated with loss of Xkr8 scramblase function can be rationalized by impaired efferocytosis and potential secondary necrosis and activation of DAMPs, the mechanisms by which tumor growth and immune activation in the TMEM16F KO E0771 cells are more elusive but nonetheless interesting. *In vitro*, the TMEM16F KO cells are impaired in calcium-mediated PS externalization suggesting that prolonged calcium signaling occurs on the viable tumor cells in the tumor microenvironment. It has long been recognized that dysregulated calcium is associated with oncogenic stress, including mitochondrial and ER stress, as well as hypoxia and glucose deprivation ^54^. We posit that one or more of these signals contribute to altered and prolonged calcium signaling in tumor cells and these cells can engage inhibitory PS receptors on immune cells in the tumor microenvironment. Indeed, very interesting studies by Wang and colleagues recently established constitutively “PS^out^” live tumor cells by knocking out the TMEM30 (CDC50A) subunit of P4 ATPases, a family of flippases that vectorially transfer PS from the outer surface to the inner surface of the plasma membrane ^66^. PS^out^ tumors developed more aggressively than wild-type (WT) tumors, showing M2 polarized tumor-associated macrophages (TAMs) and fewer tumor-antigen-specific T cells. These studies, and previous studies by Rothlin and colleagues, showing that externalized PS on live cells can inhibit TIM-3 on DCs to down-regulate co-stimulatory receptors and inhibit inflammasome signaling supports the idea of PS out on live cells is an inhibitory immune signal ^27, 67^. In the studies here, by limiting the TMEM16F-mediated PS externalization on the E0771 tumor cell, these studies support the idea that tumor cells, as a cell-intrinsic event and mechanism, can mediate in part immune evasion that occurs in the tumor microenvironment.

While our present studies are consistent with the idea of cell-intrinsic mediated events on the dying and viable tumor cells, via the cell-intrinsic loss of either Xkr8 or TMEM16F, it is still unclear what other cell types may contribute to PS externalization in the complex tumor microenvironment. For example, in the original conception of Bavituximab, a PS-targeting mAb, Thorpe and colleagues conceptualized that hypoxia and neo-vascularization promoted PS externalization on vascular endothelial cells in the tumor microenvironment, suggesting in part that Bavituximab functions as an anti-vascular agent ^32, 52, 68^. In the case for hypoxia, the recent studies by Makela and colleagues showed that populations of CD44+ cancer stem cells become PS positive and can be targeted by novel PS-directed payloads ^69, 70^. In the E0771 model here, we did not observe hypoxia mediated increased PS externalization but did not investigate a stem cell pool (SFig 2F). Further studies are also needed to address effects of hypoxia and glucose deprivation in the tumor microenvironment on PS externalization on immune cells such as macrophages, cytotoxic T cells, central memory cells, and T regulatory cells that can lead to immune exhaustion, and whether PS externalization can feedback and inhibit DCs to suppress co-stimulatory molecules and antigen presentation as alluded to above. Clearly, while the tumor-centric approach to knockout scramblases provides proof-of-concept that PS externalization on the tumor cells can drive immune-evasion, further studies using CITE-seq to identify the subsets and cell types that contribute to PS in the tumor environment are clearly warranted and meritorious.

Another emerging area of PS biology that deserves further attention is whether tumor cells up-regulate or alter expression of Xkr8 or TMEM16F as a cell intrinsic or driver event in host immune evasion and tumor progression. For example, surveying the Kaplan meier database suggest that expression levels of Xkr8 or TMEM16F are associated with poorer overall survival in breast cancer (SFig 1). Clearly, identification of signaling pathways that lead to activation and/or upregulation of Xkr8 and TMEM16F might lead to “oncogenic hubs” that link signal transduction to PS-mediated immune escape. As noted above, Xkr8 up-regulation has been observed in solid tumors following chemotherapy and radiation therapy, possibly as an adaption to oncogenic stress ^62, 66^. Moreover, phosphorylation at the carboxyl termini region of Xkr8 has been shown to activate the scramblase activity, potentially linking oncogenic kinases with PS externalization ^71, 72^. Similarly, oncogenic pathways that activation phospholipase C are long known to increase cellular IP3 and increase cytosolic calcium, potentially linking tyrosine kinase signaling to immune evasion. Finally, in recent years, emerging evidence suggests that activating mutations in flippases and scramblases can constitutively inactivate or activate enzymatic activity, possible as an oncogenic event that leads to intrinsic host immune evasion ^51^. Further studies are needed to assess the frequency of oncogenic mutations that lead to PS externalization on tumor cells in this emerging research field.

While the present study shows proof-of-concept that functional loss of function of the caspase-activated and calcium-activated scramblases and impinge on host immunity, many questions remain about the physiological context of how these scramblases are typically regulated in the tumor microenvironment. However, these studies support that further development of scramblase modifying modalities as a new line of therapeutic intervention that may have benefit in immune-oncology.

## Supplemental Figures

(S.1.A) Kaplan-Meier survival analysis of breast cancer patient data, showing a negative correlation between Xkr8 gene expression and patient survival, with higher Xkr8 levels associated with poorer survival. (1.B) Kaplan-Meier survival analysis of breast cancer patients, comparing low and high TMEM16F gene expression. Patients with low TMEM16F expression exhibited better survival compared to those with high TMEM16F expression.

(S.2.A.) Surveyor assay showing a successful KO of Xkr8 gene in EO771 cells. (S.2.B.) p-Akt signaling upon treatment of scramblase KO cells with Gas6 shows no significant difference in their intrinsic Axl-Gas6 signaling capabilities. (S.2.C.) Quantification of metastatic nodules in the lung from NSG mice injected with scramblase KO tumors. (S.2.D.) Incucyte imaging quantifying cell death upon different treatments with the Cytotox dye shows that glucose deprivation and MG132 are not toxic to cells at earlier time points. (S.2.E.) Quantification of intracellular calcium and annexin V staining on EO771 cells treated with calcium ionophore shows an early spike with calcium which subsides with time. (S.2.F.) Hypoxic conditions for 24 hours with 2% O_2_ did not induce live cell PS externalization in EO771 cells.

## Acknowledgements

We thank members of our laboratory for helpful discussions. This work was supported in part by NIH R01 CA260137-01A1 from the National Cancer Institute to RBB and SVK. Biorender was used to create diagrams. RBB and SVK disclose they are cofounders of a biotechnology company called Targeron Therapeutics that aims to develop PS-targeting IFNs for anti-cancer and anti-viral applications.

