## supplemental figures for "Phospholipid Scramblases TMEM16F and Xkr8 mediate distinct features of Phosphatidylserine (PS) externalization and immune suppression to promote tumor growth"

### Slide 1
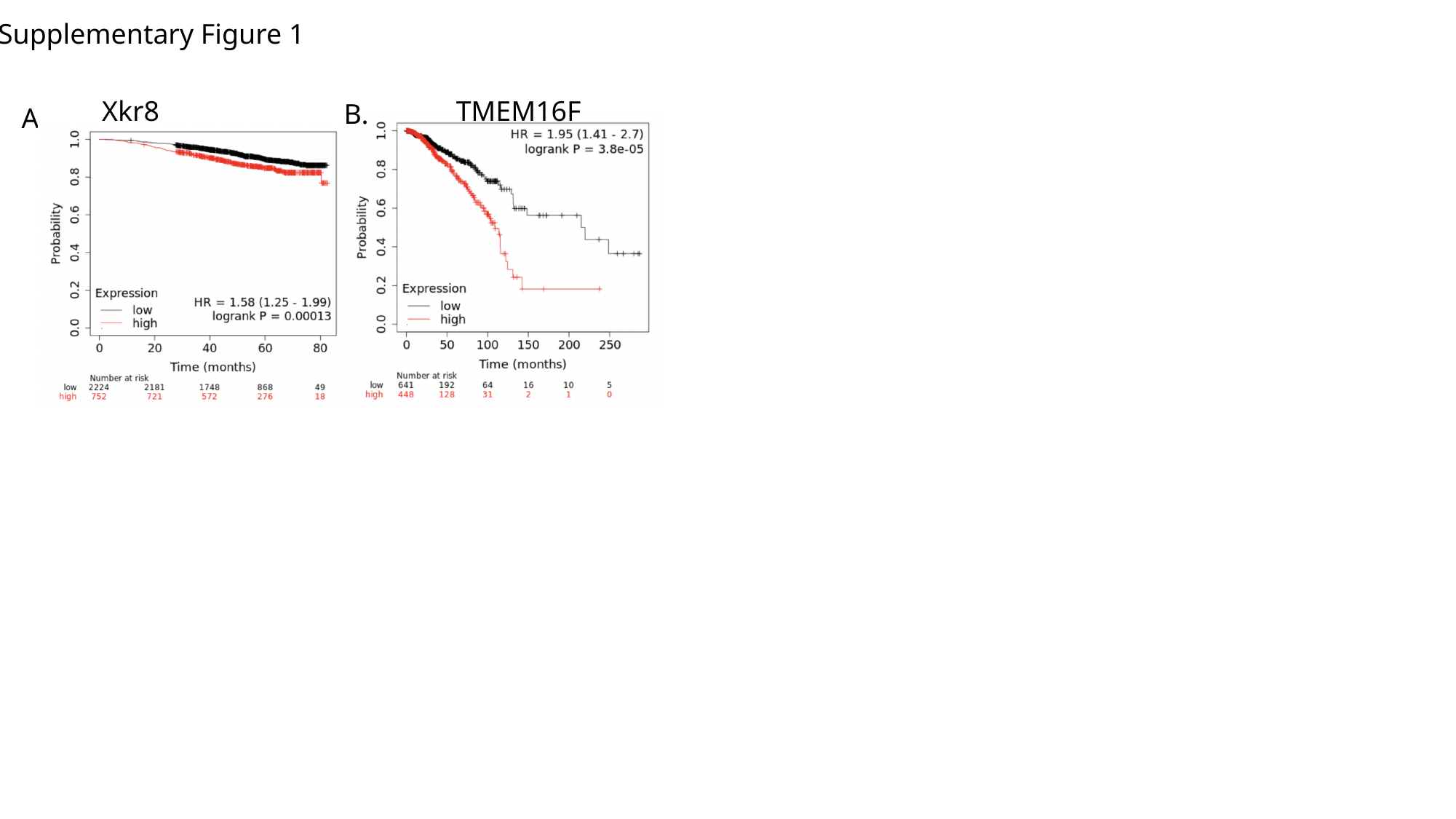

Supplementary Figure 1
Xkr8
TMEM16F
B.
A.

### Slide 2
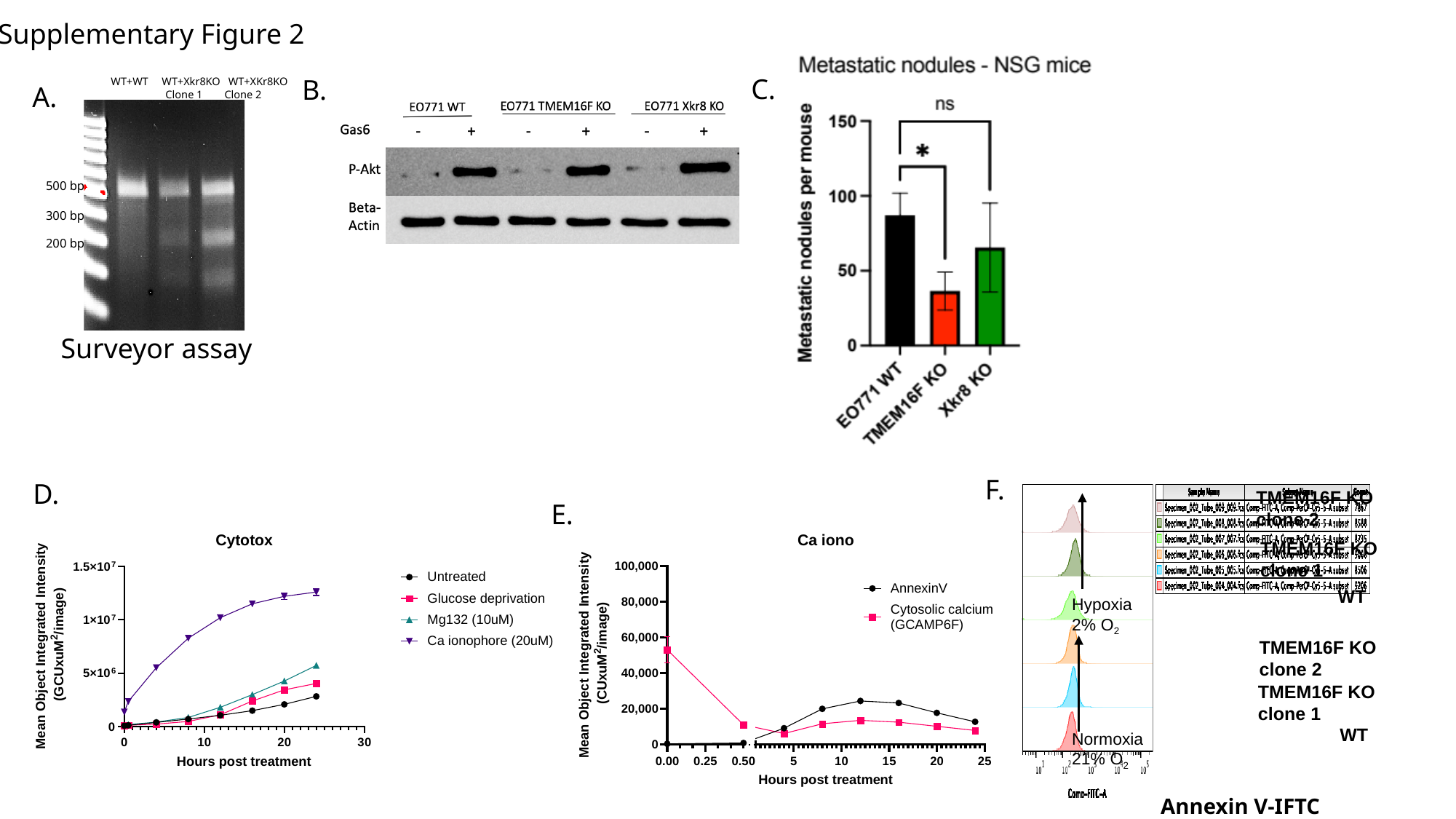

Supplementary Figure 2
C.
B.
WT+WT WT+Xkr8KO WT+XKr8KO
 Clone 1 Clone 2
500 bp
300 bp
200 bp
A.
Surveyor assay
F.
D.
TMEM16F KO clone 2
TMEM16F KO clone 1
WT
Hypoxia 2% O_2_
TMEM16F KO clone 2
TMEM16F KO clone 1
WT
Normoxia 21% O_2_
Annexin V-IFTC
E.
